# MiR-137 and miR-122, two outer subventricular zone-enriched non-coding RNAs, regulate basal progenitor expansion and neuronal differentiation

**DOI:** 10.1101/2021.04.01.438039

**Authors:** Ugo Tomasello, Esther Klingler, Mathieu Niquille, Nandkishor Mule, Laura de Vevey, Julien Prados, Antonio J Santinha, Randall Platt, Victor Borrell, Denis Jabaudon, Alexandre Dayer

## Abstract

Cortical expansion in the primate brain relies on the presence and the spatial enlargement of multiple germinal zones during development and on a prolonged developmental period. In contrast to other mammals, which have two cortical germinal zones, the ventricular zone (VZ) and subventricular zone (SVZ), gyrencephalic species display an additional germinal zone, the outer subventricular zone (OSVZ), which role is to increase the number and types of neurons generated during corticogenesis. How the OSVZ emerged during evolution is poorly understood but recent studies suggest a role for non-coding RNAs, which allow tight regulations of transcriptional programs in time and space during development (Dehay et al. 2015; Arcila et al., 2014). Here, using *in vivo* functional genetics, single-cell RNA sequencing, live imaging and electrophysiology to assess progenitor and neuronal properties in mice, we identify two ferret and human OSVZ-enriched microRNAs (miR), miR-137 and miR-122, which regulate key cellular features associated with cortical expansion. MiR-137 promotes basal progenitor self-replication and superficial layer neuron fate, while miR-122 slows down neuronal differentiation pace. Together, these findings support a cell-type specific role for miR-mediated transcriptional regulation in cortical expansion.

## Introduction

While the human cortex has circonvolutions, many species, including within primates, have a smooth cortex instead (Rakic, 2000, 2009). Adult cortical neuron numbers depend in particular on the number of progenitors present during corticogenesis, their ability to self-amplify, and the duration of the neurogenic period. In mammals, there are two main types of cortical progenitors, which reside in distinct germinal zones: (1) the ventricular zone (VZ) hosting apical progenitors (APs, also called apical radial glia) (Malatesta et al., 2000; Noctor et al., 2001) abuts the ventricular wall, and (2) the subventricular zone (SVZ), located more superficially, which contains basal progenitors (BPs), that are transit amplifying cells expanding neuronal output from APs (Noctor et al., 2004; Haubensak et al., 2004; Miyata et al., 2004). In gyrencephalic species, BPs are thought to have undergone an increase in their proliferative capacity, resulting in a thickening of the SVZ and emergence of anatomical subcompartments in the form of an “inner” SVZ (iSVZ) and “outer” SVZ (oSVZ) (Reillo et al., 2012). This expansion is particularly visible at later stages of corticogenesis, as superficial layer neurons are being produced.

How SVZ expansion occurred during evolution is still poorly understood, but one possibility is the implementation of fine regulatory transcriptional controls over the proliferation and differentiation of progenitors and neurons (Fietz et al., 2010, 2012; Reillo et al., 2011, 2012). Non-coding RNAs and, especially, microRNAs are particularly interesting candidates in this respect as their numbers increase with evolution (Liu et al., 2013). Non-coding RNAs fine-tune various aspects of neurogenesis across species, allowing BP amplification in the oSVZ and, secondarily, cortical expansion, in particular through formation of gyri (Martinez-Martinez et al., 2016, Nowakowski et al., 2018, Arcila et al., 2014, Ayoub et al., 2011, Fietz at al., 2012, de Juan Romero et al., 2015, Diaz et al., 2020). Supporting a role for microRNAs in cortical expansion, here we show that overexpression of the ferret and human oSVZ-enriched microRNAs miR-137 and miR-122 in the mouse neocortex, which is normally devoid of these microRNAs, leads to increased BP self-replication and slowing down of neuronal differentiation pace, respectively. miR-137 acts in progenitors, expanding BP numbers by increasing proliferative divisions, ultimately resulting in increased superficial layer neuron production, whereas miR-122 acts in newborn neurons by slowing their differentiation pace, potentially allowing longer distances to be covered before differentiation occurs. Together, these results reveal a cell-type specific role for microRNAs in the regulation of cortical expansion.

## Results

### MiR-122 and miR-137 are enriched in the ferret cortex oSVZ during SL neurogenesis

To identify microRNAs that could regulate neurogenic programs in gyrencephalic species, we first performed a screen using existing RNA expression data obtained from the three germinal compartments of the developing gyrencephalic ferret cortex (VZ, iSVZ and oSVZ) at postnatal day (P) 2, when superficial layer (SL) neurons are being born (de Juan Romero et al., 2015). This analysis revealed 23 oSVZ-enriched microRNAs, with miR-137 and miR-122 being the most strongly differentially expressed (**Fig. 1A, bottom right**). This enrichment was specific to the period of generation of SL neurons, since analysis at earlier time points (embryonic day (E)30, at the beginning of cortical neurogenesis, and E34, when deep layer (DL) neurons are being born) did not reveal a similar enrichment (**Fig. 1A** **bottom left**, Martinez-Martinez et al., 2016). MiR-137 is also enriched in the embryonic oSVZ of macaques (Arcila et al., 2014), and in humans, both miR-137 and miR-122 are enriched in germinal zones during SL neurogenesis (Nowakowski et al., 2018, **Fig. S1A,B**). Consistent with the lack of a distinctive oSVZ compartment in the mouse, only very low levels of both miRs were detected at E15.5 using a high-sensitivity double sensor tool (DFRS), in which the target sequence of the microRNA controls the expression of a modified RFP (De Pietri Tonelli et al., 2006, **Fig. S1D** and see Methods), while they were not even detectable using *in situ* hybridizations (**Fig. S1E**), consistent with previous data using microarray during mouse SL neurogenesis (Volvert et al., 2014).

**Figure 1:**
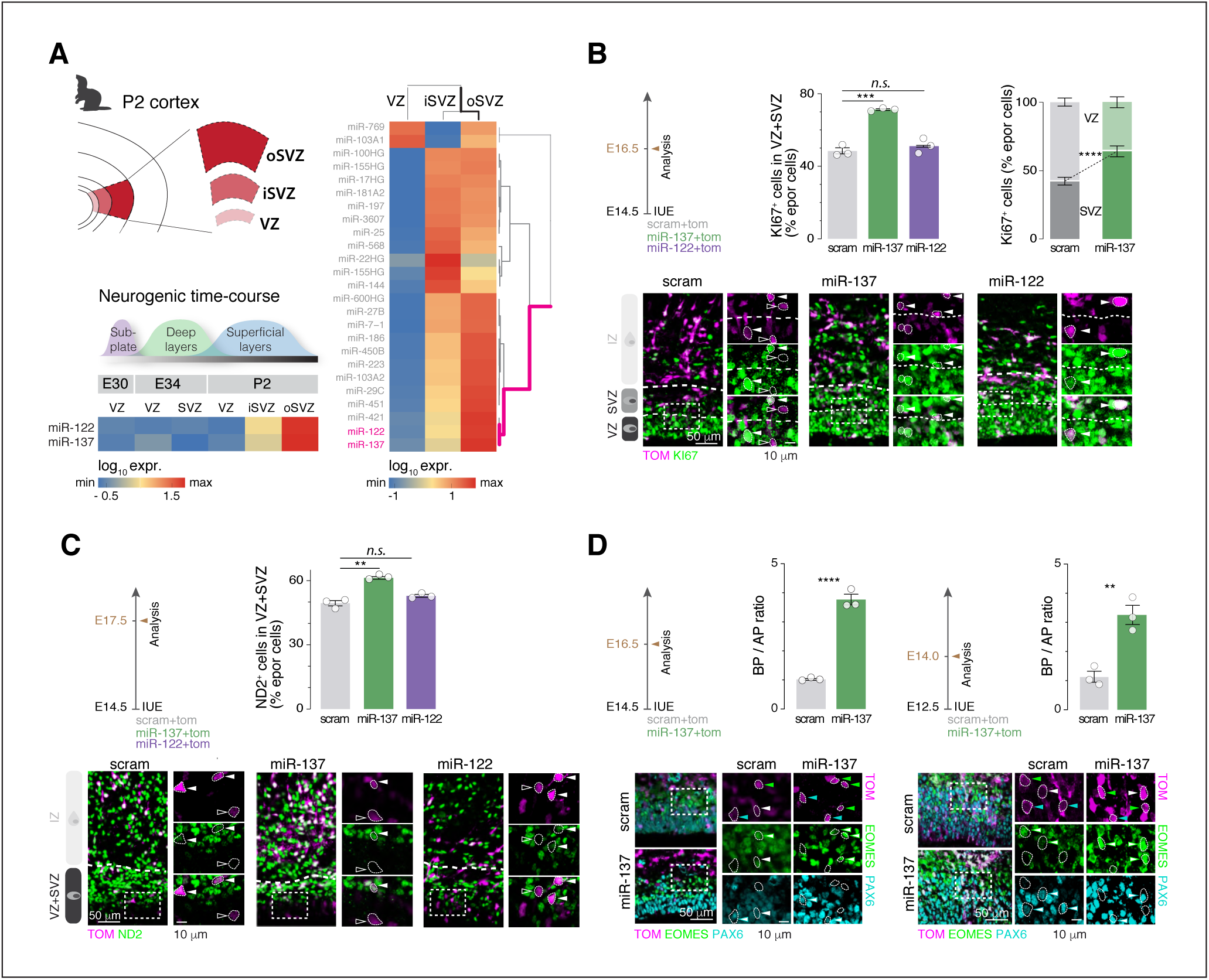
oSVZ enriched MiR-137 but not miR-122 affects cortical progenitor proliferation. (**A**) Microarray of ferret ventricular (VZ), inner and outer subventricular (iSVZ and oSVZ, respectively) zones during cortex development (data are from de Juan Romero et al., 2015 and Martinez-Martinez et al., 2016 for postnatal day (P) 2 and embryonic day (E) 30 / E34 / P2, respectively). Top left, schematic of microdissections of P2 cortex performed for the collections of P2 VZ, iSVZ and oSVZ. Right, differentially expressed microRNAs in the three germinal zones at P2. Note that miR-122 and miR-137 cluster together as the most enriched in the oSVZ. Bottom left, Expression of miR-137 and miR-122 in VZ and SVZ along development. (**B**) Progenitors in the VZ and SVZ at E16.5 upon miR-137 or miR-122 overexpression in E14.5 mouse cortex. Top left, experimental design. Top right, quantifications of KI67+ electroporated cells in VZ and SVZ. Bottom, representative micrographs of KI67+ electroporated cells. Electroporated cells positive (filled arrowheads) and negative (empty arrowheads) for KI67 are highlighted. (**C**) Neurogenic output at E17.5 upon miR-137 or miR-122 overexpression in E14.5 mouse cortex. Top left, experimental design. Top right, quantifications of NeuroD2 (ND2)+ electroporated cells in VZ and SVZ. Bottom, representative micrographs of electroporated cells labeled for ND2. Electroporated cells positive (filled arrowheads) and negative (empty arrowheads) for ND2 are highlighted. (**D**) Basal progenitor (BP) *versus* apical progenitor (AP) ratio upon miR-137 overexpression at E14.5 (left) or E12.5 (right) addressed using immunohistochemistry against EOMES to identify BP and PAX6 to identify AP. Bottom, representative micrographs of EOMES / PAX6 labelings. Green arrowheads, EOMES^+^ electroporated cells; cyan arrowheads, PAX6+ electroporated cells, white arrowheads, EOMES^+^/PAX6^+^. Data are represented as mean ± SEM. Biological replicates are distinguished by circles in the bar plots. One-way ANOVA (B, bottom right; C; D, left); Two-way ANOVA (B, bottom left); Unpaired t-test (D, right). *p < 0.05, **p < 10^-2^, ***p < 10^-3^, ****p < 10^-4^.

### MiR-137 but not miR-122 affects cortical progenitor proliferation in mice

Based on the spatial and temporal specificity of expression of miR-137 and miR-122, we hypothesized that these microRNAs could regulate salient properties of oSVZ progenitors and their progenies. To test this hypothesis, we used a gain-of-function approach in mice, in which oSVZ-type progenitors are very rare (Shitamukai et al., 2011; Wang et al., 2011); we used *in utero* electroporation (IUE) to overexpress miR-137 or miR-122 at embryonic day (E)14.5, the time when SL neuron generation is starting (**Fig. S1E** and see Methods). We first performed a time-course analysis 24, 48 and 72 hours (hrs) after electroporation (*i.e.* at E15.5, E16.5 and E17.5) to assess the effect of miR-137 and miR-122 on progenitor proliferation (using KI67 as marker for cycling cells). We found that miR-137 expanded progenitor numbers (KI67^+^ cells) 48 hrs after IUE (E16.5, **Fig. 1B**). Most of these progenitors were located in SVZ, where BPs normally reside (**Fig. 1B** **top right**). In contrast, miR-122 did not affect KI67^+^ cell numbers (**Fig. 1B**). We also followed the fate of miR-137 and miR-122-overexpressing cells using NeuroD2 immunostaining to identify post-mitotic neurons (**Fig. 1C**). This analysis revealed that while miR-137 increased neuronal production 72 hrs after IUE (E17.5), miR-122 had no such effect (**Fig. 1C**). Using progenitor identity markers PAX6 and EOMES to distinguish APs and BPs by immunostaining, we found a 3-fold increase in BP / AP ratio following miR-137 expression (**Fig. 1D**). A similar increase in BP numbers was also observed following E12.5 electroporation, suggesting that the transcriptional networks modulated by miR-137 are accessible before SL neuron generation (**Fig. 1D, right**). Together, these results reveal that miR-137 overexpression increases basal progenitor pool and subsequent neuron numbers in the mouse neocortex.

### MiR-137 overexpression promotes basal progenitor generation and proliferation

Focusing on miR-137, we sought to identify cell-type specific transcriptional programs that are regulated by this microRNA by performing single cell-RNA sequencing 24 hrs following its expression. A scrambled (scram) version of the miR-137 was used in control experiments. We distinguished APs, BPs, newborn (N_0_) and maturing neurons (N_1_) based on their transcriptional identity (Telley, Agirman et. al, 2019) (**Fig. 2A**). As expected and confirming the role of miR-137 in the expansion of BP pool, BPs were in higher proportions amongst progenitors in the miR-137 compared to scrambled conditions (**Fig. 2B**). We designed a support vector machine learning approach to identify genes induced and repressed in each cell type, which revealed cell-type specificity in the genes regulated by miR-137 (**Fig. 2C**, **left**): BPs were most affected, followed by N_0_ and AP (**Fig. 2C**, **right**). Focusing on BPs, we next analyzed the most strongly repressed and induced genes in this cell type: repressed genes were mostly associated with mitochondrial activity and neurogenic processes while induced genes were related to cell cycle progression (**Fig. S2A**). Using gene expression data across corticogenesis from our lab (Telley et al., 2019), we showed that miR-137 induced genes in BPs were mainly expressed in progenitors, while miR-137 repressed genes were expressed in neurons (**Fig. S2B**). Consistently, control neurons from our dataset showed higher expression of miR-137 repressed genes in BPs than control APs (**Fig. S2C**). Together, these results suggest that miR-137 promotes proliferative and represses neuronal properties of BPs. Interestingly, human basal progenitor cells also retain high levels of proliferative genes compared to APs (**Fig. S2D**), suggesting a potential evolutionary relevance for the effects observed with miR-137 overexpression in mice. As a proof-of-principle for the functional genes regulated by miR-137 in BPs, we functionally investigated two targets, *Cd63* (amongst the most induced genes, **Fig. S2A**) and *Myt1l* (amongst the most repressed genes, **Fig. S2A**), which encode an extracellular matrix receptor (Tschoepe et al., 1990) and a transcription factor critical for neuronal identity (Mall et al., 2017), respectively. Consistent with a role in proliferation, gene overexpression led to an increase (by *Cd63*) or a decrease (by *Myt1l*) in progenitor number and BP / AP ratio (**Fig. 2D**, **Fig. S2E**). Together, these results indicate that miR-137 regulates key transcriptional programs to promote BP generation and proliferation allowing the dramatic expansion of superficial layers characterizing more evolved species (**Fig. 2E**).

**Figure 2:**
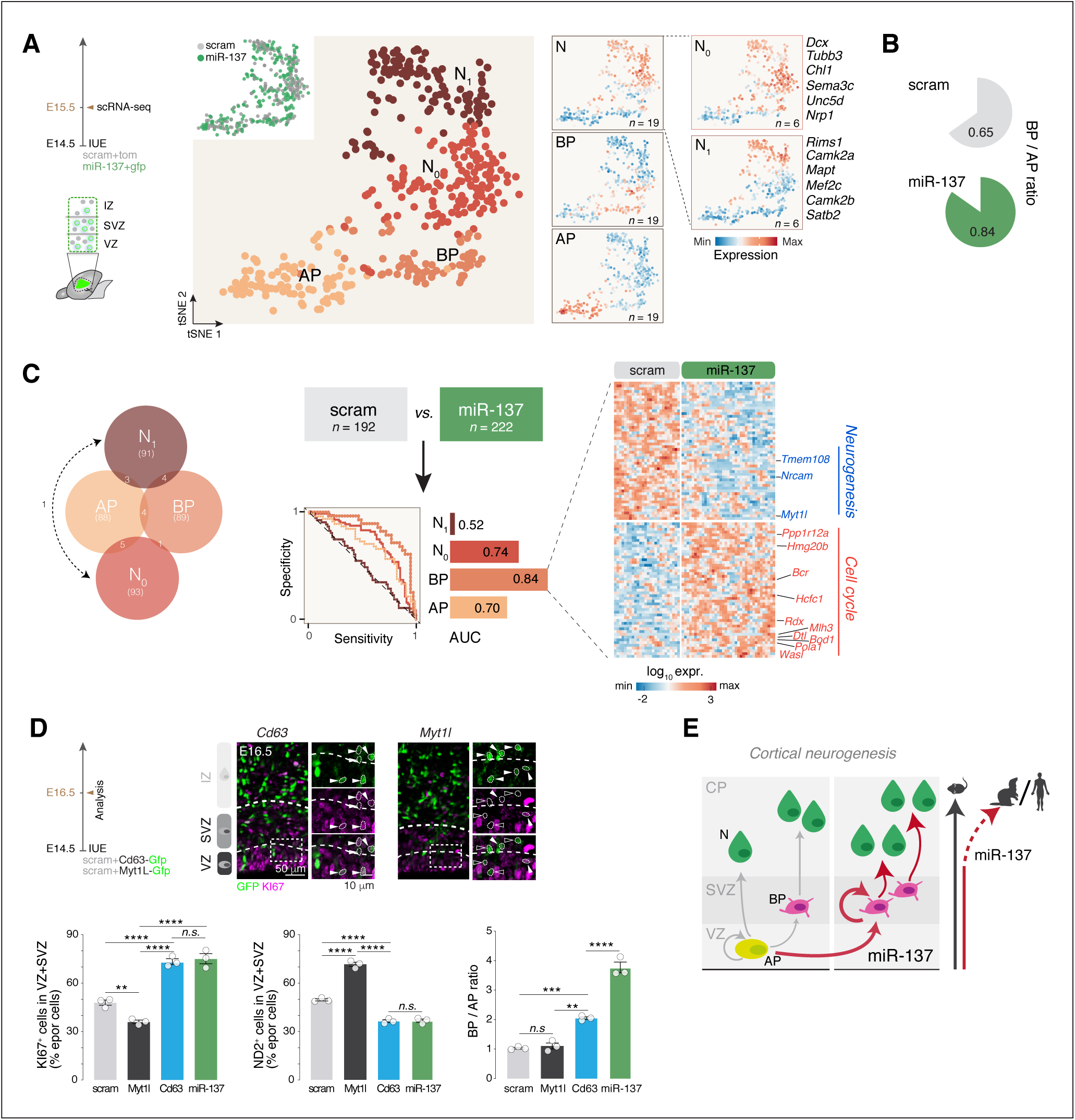
miR-137 regulates transcriptional programs promoting BP generation and proliferation. (**A**) T-SNE representation of E15.5 scramble (scram) and miR-137 E14.5-*in utero* electroporated (IUE) scRNA-seq data reveals transcriptional organization of the cells according to their differentiation status. Apical progenitors (AP), basal progenitors (BP), newborn neurons (N_0_) and differentiating neurons (N_1_) can be distinguished by their combinatorial expression of key marker genes. (**B**) BP / AP ratio in scram and miR-137 conditions. (**C**) Machine learning approach to identify cell type-specific core sets of genes classifying neurons and progenitors in miR-137 and scram conditions. Left, genes differentially expressed shared between cell types. Middle, Machine learning prediction score for each cell type showed as the area under the specificity / sensitivity curves (AUC). Note the highest score in BP. Right, Heatmap of top miR-137 induced and repressed genes in BP. (**D**) Progenitor and neurons number and BP / AP ratio upon overexpression of *Cd63* (induced by miR-137) or *Myt1l* (repressed by miR-137) at E16.5. Progenitors were identified as KI67+ cells, neurons as NeuroD2 (ND2)^+^ cells, BP and AP as EOMES^+^ and PAX6^+^ cells, respectively. Top, representative micrographs of KI67 immunohistochemistry upon *Cd63* or *Myt1l* overexpression. (**E**) Schematic representation of basal progenitor increase through self-amplification and role of miR-137 in SL neurogenesis and expansion across evolution. Data are represented as mean ± SEM. Biological replicates are distinguished by circles in the bar plots. One-way ANOVA (D). *p < 0.05, **p < 10^-2^, ***p < 10^-3^, ****p < 10^-4^.

### MiR-137 overexpression promotes SL-type neuron fate

We next examined the fate of neurons generated following miR-137 overexpression. During normal corticogenesis, the number of BPs and size of the SVZ increase, such that a significant fraction of SL neurons are born from divisions of BPs, which contrasts with earlier stages of corticogenesis (Noctor et al., 2004, 2008; Kowalczyk et al., 2009; Taverna et al., 2014). We thus hypothesized that the expansion of the BP numbers by miR-137 would promote L2/3 neuron fate. To investigate this possibility, we first examined the laminar distribution of neurons born following miR-137 IUE at E14.5, the time of birth of L4 neurons. We performed this analysis at P7, a time at which neuronal migration is essentially complete. In contrast to the control condition, in which neurons were predominantly located in L4, neurons in the E14.5 miR-137 overexpression condition were located more superficially, including within L2/3 (**Fig. 3A**). We next asked whether this increase in L2/3 neurons was due to an increase of late neurogenesis and / or to a shift in L4-to-L2/3 identity. We first chronically applied EdU to birth-label neurons as BPs are dividing (E16.5, **Fig. 3B**). The fraction of neurons labelled in the miR-137 overexpression condition was 3 times higher than in the control condition (30% *vs*. 10%) suggesting that miR-137 amplifies neuronal production increasing the L2/3 neuron output (**Fig. 3B**). In order to address the molecular L4 *vs*. L2/3 identity at P7 upon miR-137 overexpression, we performed scRNA-seq of P7 neurons after E14.5 electroporation of scrambled or miR-137 (**Fig. 3C**). We used a control dataset of P7 neurons electroporated with scrambled at E14.0 (putative L4 neurons) and E15.5 (putative L2/3 neurons) to predict the molecular identities of miR-137-overexpressing neurons at P7 using a machine-learning classifier (**Fig. 3C**, **Fig. S3A**). This approach showed an increased proportion of neurons with L2/3-type identity in the miR-137 overexpression condition, suggesting acquisition of L2/3 neuron-type identity (**Fig. 3C**). Supporting this finding, L4-located neurons in the E14.5 miR-137 condition did not express the L4 neuron marker RORB (**Fig. 3D, Fig. S3B**). Consistent with a functional relevance of this L4-to-L2/3 shift in molecular programs, L4 neurons in the E14.5 miR-137 overexpression condition often displayed an apical dendrite (which is typically absent in L4 neurons and instead is present in L2/3 neurons, **Fig. 3E**) and displayed an Ih current, an electrophysiological feature normally not found in L4 neurons and instead present in L2/3 neurons (**Fig. 3F**; De La Rossa et al., 2013). Finally, retrograde labeling from the contralateral hemisphere revealed the presence of interhemispheric projection in L4-located neurons following miR-137-overexpression at E14.5, a typical feature of L2/3-type neurons (**Fig. 3G**). Overall, these results indicate that the amplification of the BP pool by miR-137 results in the expansion of neurons with laminar, molecular, anatomical and functional features of L2/3-type neurons (**Fig. 3H**).

**Figure 3:**
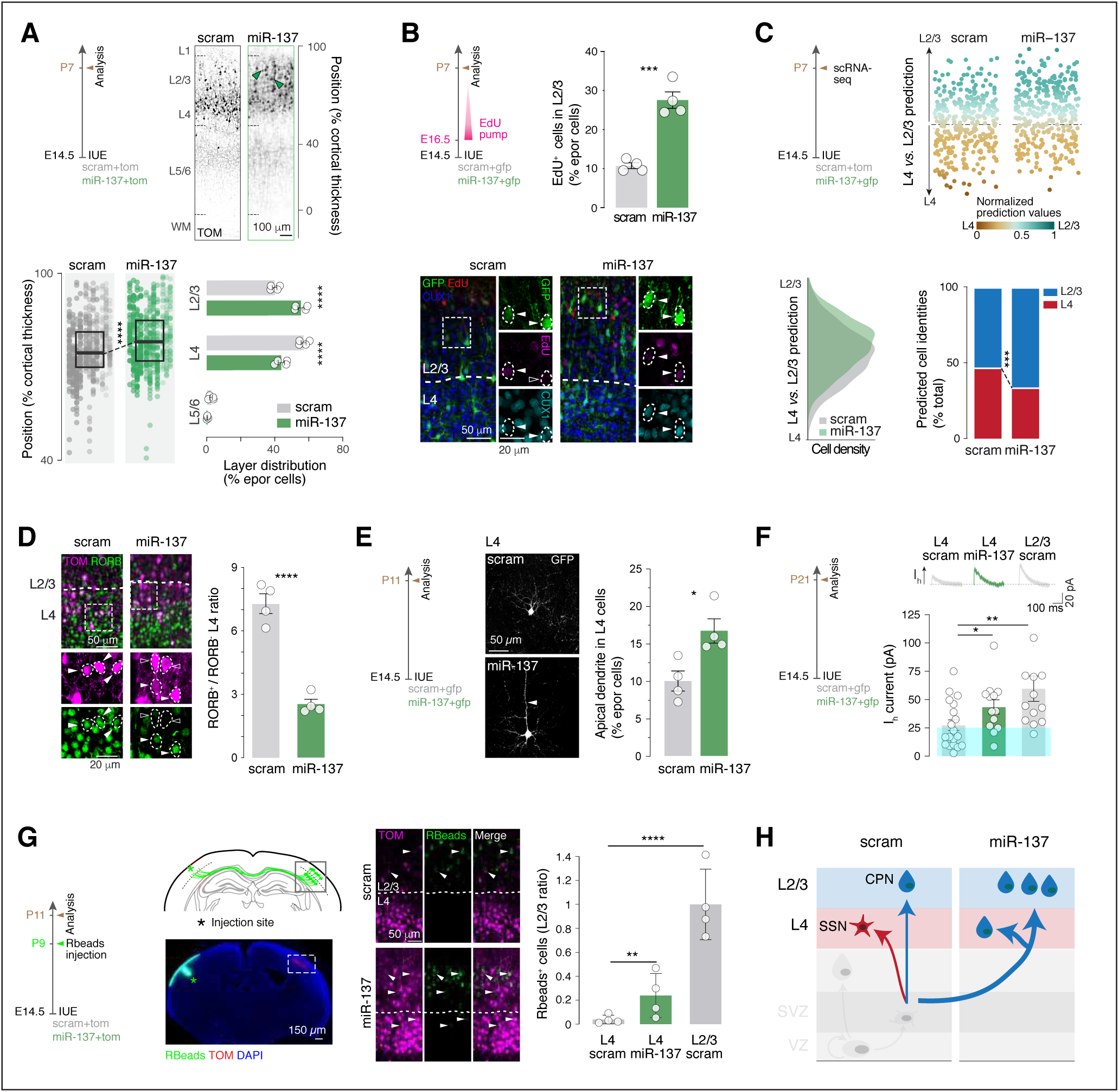
miR-137 promotes the expansion of L2/3 by increasing late neurogenesis and reprogramming L4 neurons. (**A**) Radial position of neurons electroporated (epor) with scramble (scram) or miR-137 at P7. Note the shift of position towards L2/3 in miR-137 condition. (**B**) EdU labeling of neurons born after E16.5 upon miR-137 overexpression. Bottom, representative micrographs of EdU staining at P7 in superficial layers labeled with CUX1. Note that miR-137 increases late neurogenesis. Electroporated cells positive (filled arrowheads) and negative (empty arrowheads) for EdU are highlighted. (**C**) ScRNA-seq of P7 scram and miR-137 neurons electroporated at E14.5 and of P7 scram neurons electroporated at E14 (L4) and E15.5 (L2/3). L4 *versus* L2/3 molecular identities of E14.5 electroporated scram and miR-137 cells were calculated using the scram L4 *versus* L2/3 prediction model (top). Bottom, prediction model cell density (left) and proportion of cells with L4 *versus* L2/3 identities (right) in scram and miR-137 conditions. (**D**) RORB expression in L4 at P7. Left, representative micrographs of immunohistochemistry against RORB. Right, quantification of RORB^+^ *versus* RORB^-^ cells in L4 electroporated (epor) cells. Electroporated cells positive (filled arrowheads) and negative (empty arrowheads) for RORB are highlighted. (**E**) Morphology of L4 neurons at P11. Left, representative micrographs of L4 scram and miR-137 IUE neurons at P11. Right, quantification of the L4 neurons with an apical dendrite (arrowhead). (**F**) I_h_ current in P21 L4 and L2/3 scram and L4 miR-137 neurons. Top right, representative traces. (**G**) Retrograde labeling of callosally-projecting neurons using retrobeads (Rbeads). Left, experimental design. Rbeads were injected in the controlateral cortex to the electroporation at P9. Right, quantifications of Rbeads+ electroporated cells in L4 and L2/3 in scram and miR-137 conditions. Data were normalized by the number of Rbeads^+^ cells in L2/3 scram condition. Arrowheads, Rbeads electroporated cells. (**H**) Schematic diagram of the role of miR-137 in L2/3 expansion. SSN, spiny stellate neurons; CPN, callosally-projecting neurons. Data are represented as mean ± SEM. Biological replicates are distinguished by circles in the bar plots. One-way ANOVA (A, bottom left); Two-way ANOVA (A, bottom right; D, G); Fisher’s chi-square test (C); Unpaired t-Test (B, E, F). *p < 0.05, **p < 10^-2^, ***p < 10^-3^, ****p < 10^-4^.

### MiR-122 regulates the unfolding of postmitotic neuronal differentiation programs

As indicated above miR-122 does not affect progenitor proliferation (**Fig. 1B-C**), but still could act on the differentiation of newborn neurons in the SVZ during late neurogenesis. To investigate a potential postmitotic effect of miR-122, we overexpressed this transcript using IUE, as described above. Using the same approach than for miR-137, we performed single-cell RNA sequencing 72 hrs following miR-122 expression, distinguishing APs, BPs, newborn (N_0_) and maturing neurons (N_1_) based on their transcriptional identity (**Fig. 4A**). Support vector machine learning approach identified induced and repressed cell type-specific pools of genes regulated by miR-122 (**Fig. 4B**). Consistent with a postmitotic effect, the analysis revealed that maturing neurons (N_1_) were the most affected by miR-122 (**Fig. 4C**). MiR-122 target genes within these neurons had ontologies relating to cell morphogenesis and differentiation, including microtubule cytoskeleton structure and axon development (**Fig. 4C** **and Fig. S4A-B**). Consistent with a role in neuronal maturation, pseudotime analysis revealed that miR-122 overexpressing neurons were transcriptionally less mature than their control counterparts, suggesting that miR-122 overexpression delayed neuronal maturation (**Fig. 4D**). To assess the functional correlate of these transcriptional changes, we analyzed key neuronal post-mitotic features such as positioning, migration dynamics and expression of mature molecular programs in miR-122-overexpressing neurons. This first revealed that miR-122 overexpressing neurons were mostly found between the SVZ and the lower IZ, while only few cells reached the cortical plate (**Fig. 4E**). Using live imaging in acute cortical slices, we found that the migration of miR-122-overexpressing neurons was delayed compared to that of control neurons (**Fig. 4F**). Consistent with a slower maturation process, miR-122 newborn neurons also failed to hyperperpolarize their membrane potential to levels normally found in control neurons (**Fig. 4G**; Picken Bahrey et al., 2003). Further supporting slower differentiation, SATB2 (a marker of mature SL neurons) immunofluorescence levels were lower in miR-122 overexpressing neurons than in controls (**Fig. S4C**). Finally, we compared mouse and human embryonic neurons during SL neurogenesis (Nowakowski et al., 2018, GSW 18-22 mature neurons, which we compared to our N_1_ type). Projection of human neurons in the mouse pseudo-maturation axis showed that they were even more immature than miR-122 overexpressing neurons (**Fig. 4H, Fig. S4D**), accordingly with longer differentiation times in human (Linaro et al., 2019). Altogether, these results suggest a post-mitotic role for miR-122 in slowing the pace of neuronal differentiation across evolution.

**Figure 4:**
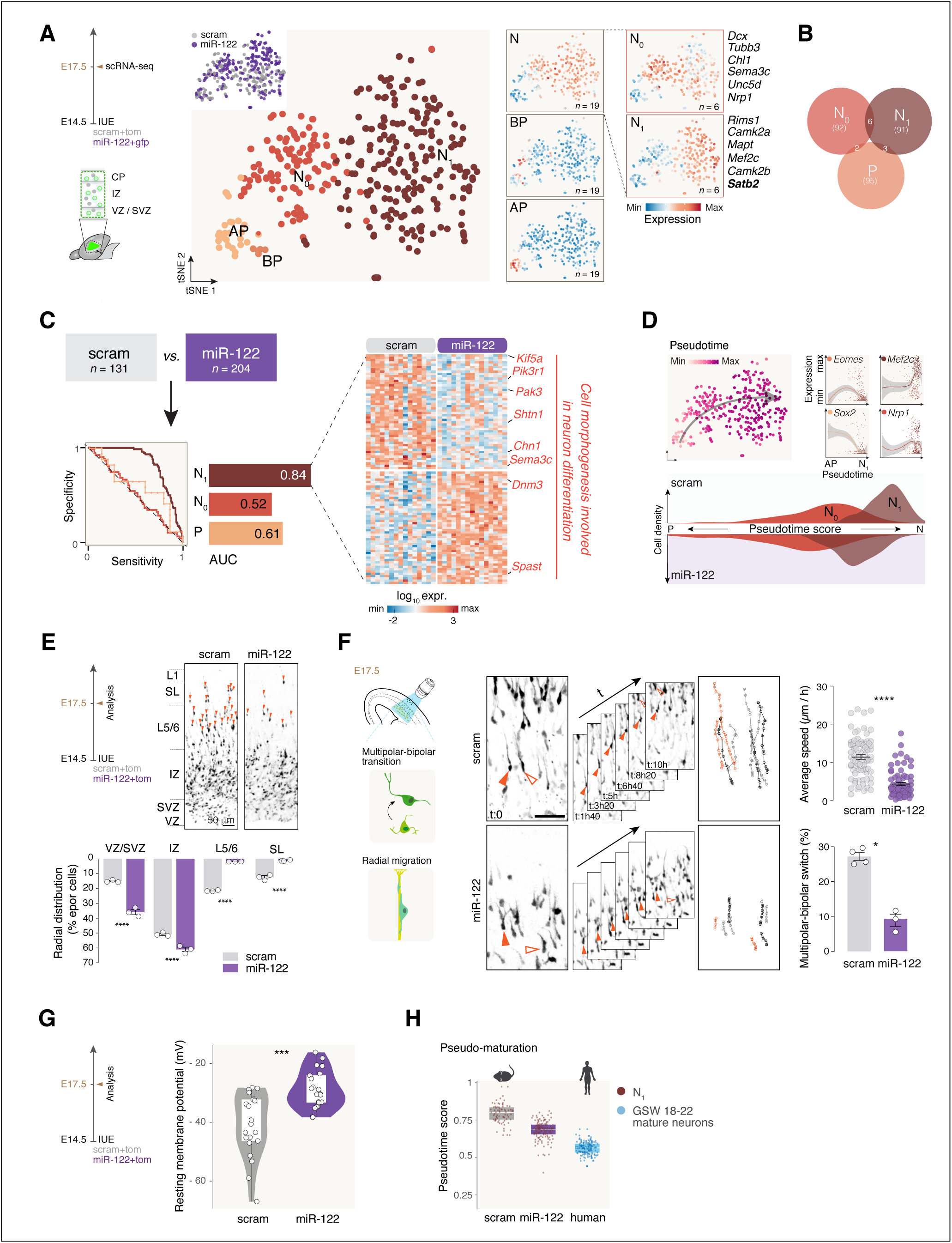
miR-122 regulates the unfolding of postmitotic neuronal differentiation programs. (**A**) T-SNE representation of E17.5 scramble (scram) and miR-122 E14.5 *in utero* electroporated (IUE) scRNAseq data reveals transcriptional organization of the cells according to their differentiation status. Apical progenitors (AP), basal progenitors (BP), newborn neurons (N_0_) and differentiating neurons (N_1_) can be distinguished by their combinatorial expression of key marker genes. (**B**) Machine learning approach to identify cell type-specific core sets of genes classifying neurons and progenitors in miR-122 and scram conditions. Here are showed shared differentially expressed genes between cell types. (**C**) Left, Machine learning prediction score for each cell type showed as the area under the specificity / sensitivity curves (AUC). Note the highest score in N1. Right, Heatmap of top miR-122 induced and repressed genes in N1. (**D**) Pseudotime analysis of scram and miR-122 single cells. Top left, Pseudotime values of single cells showed in the t-SNE space. Top right, marker genes of AP, BP, N0 and N1 expressed along pseudotime. Bottom, Density plot of pseudotime values of N0 and N1 in scram and miR-122 conditions. (**E**) Radial position of SL neurons at E17.5 after scram and miR-122 IUE. Orange arrowheads display neurons in CP. (**F**) Live imaging of scram and miR-122 IUE neurons at E17.5. Left, Experimental design. Middle, Illustrative images of neuron movement tracking in scram and miR-122 conditions. Right, quantifications of the average speed of migration and the multipolar-bipolar transition (MBT). (**G**) Resting membrane potential of scram and miR-122 migrating neurons at E17.5. (**H**) Pseudotime value predictions of human superficial layer differentiating neurons from gestational weeks (GSW) 18-22 fetuses using the mouse model. Data are represented as mean ± SEM. Biological replicates are distinguished by circles in the bar plots. (E) Two-way ANOVA; (F) Kruskal-Wallis; (G) Unpaired t-Test. *p < 0.05, **p < 10^-2^, ***p < 10^-3^, ****p < 10^-4^. Human scRNA-seq data are from Nowakowski et al., 2017.

### MiR-122 overexpression promotes L2/3-type neuron fate

We next examined the fate of neurons generated following miR-122 overexpression. First, we examined the laminar distribution of neurons at P7 following miR-122 IUE at E14.5. In contrast to the control condition, upon miR-122 overexpression half of the neurons were mispositioned in DL and WM, while the other half settled correctly in SL (**Fig. 5A**). While L2/3 neuronal distribution was similar to that of controls, L4 was essentially depleted of neurons in the miR-122 condition, suggesting that the majority of misplaced neurons in DL/WM were prospective L4 neurons. Chronic EdU labeling from E15.5 (when L2/3 neurons are born) showed that very few EdU^+^ miR-122 overexpressing cells were found in DL and WM at P3 (when migration of SL neurons is largely complete), further confirming that the vast majority of DL misplaced neurons were prospective L4 neurons (**Fig. S5A**). We next investigated the molecular identity of miR-122 overexpressing neurons (see **Fig. S5** and Methods) which revealed that miR-122 neurons in SL and DL displayed an increased ratio of L2/3-predicted compared to L4-predicted cells (**Fig. 5B**), suggesting that slowing down the differentiation of L4 neurons by miR-122 (**Fig. 4**) results in the acquisition of a L2/3 neuron-like molecular identity. Supporting this finding, L4-located miR-122 overexpressing neurons at P7 did not express the L4 neuron marker RORB (**Fig. 5C**). We next asked whether miR-122 SL and DL neurons still showed immaturity traits at P7. For this, we used a postnatal maturation model: neurons of L2/3 (E15.5 born) and L4 (E14.0 born) were collected at P3 and P7 (**Fig. S5C**). Analysis revealed that miR-122 neurons had a lower pseudo-maturation score than control neurons at P7, whatever their layer position, the DL neurons being the most immature (**Fig. 5D**). These results suggest that mispositioned miR-122 neurons may still be migrating to their final position, and therefore expressed more immature transcriptional programs. Consistently, at P21 few miR-122 neurons remained in the DL or WM and their distribution was shifted toward L2/3 compared to scram neurons in SL (**Fig. 5E**, **Fig. S5D**). Since we did not observe increased cell death in miR-122 neurons (**Fig. S5E**) and in line with the shift in neuronal identity reported above, these results suggest a protracted migration upon miR-122 overexpression and a preferential settling in L2/3. Overall, these results reveal that miR-122 affects the maturation pace of SL neurons and suggest that slowing of transcriptional programs by miR-122 overexpressing L4-type neurons is associated with acquisition of L2/3-type identity (**Fig. 5F**).

**Figure 5:**
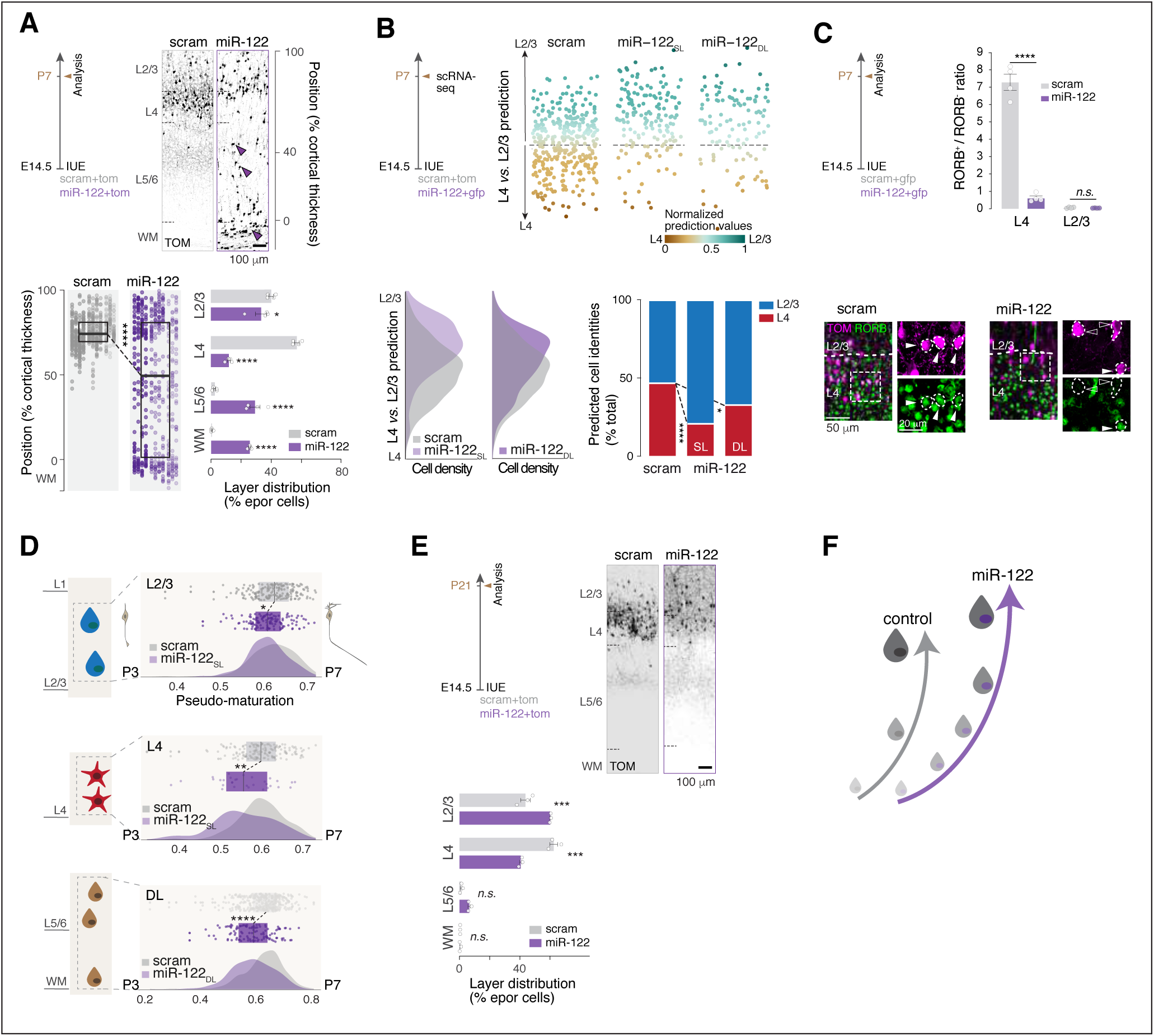
miR-122 overexpression slows the pace of neuronal differentiation and promotes L2/3-type neuron fate. (**A**) Radial position of neurons electroporated (epor) with scramble (scram) or miR-122 at P7. Arrowheads, neurons in deep layers or white matter (WM). (**B**) ScRNA-seq of P7 scram and miR-122 neurons electroporated at E14.5 and of P7 scram neurons electroporated at E14 (L4) and E15.5 (L2/3). L4 *versus* L2/3 molecular identities of E14.5 electroporated scram and miR-122 cells were calculated using the scram L4 *versus* L2/3 prediction model (top). Bottom, prediction model cell density (left) and proportion of cells with L4 *versus* L2/3 identities (right) in scram and miR-122 conditions. miR-122 cells predicted as positioned in superficial (SL) or deep (DL) layers are represented apart. (**C**) RORB expression in L4 at P7. Top, quantification of RORB^+^ *versus* RORB^-^ cells in L4 epor cells. Bottom, representative micrographs of immunohistochemistry against RORB. Electroporated cells positive (filled arrowheads) and negative (empty arrowheads) for RORB are highlighted. (**D**) Pseudo-maturation score calculated using a model of P3 *versus* P7 L4 and L2/3 neurons (electroporated with scram at E14 and E15.5, respectively). L2/3 and L4 neurons are represented apart. miR-122 neurons located in DL were compared to all scram SL neurons (light grey). (**E**) Radial position of neurons epor with scram or miR-122 at P21. Note the absence of neurons in DL in the miR-122 condition. (**F**) Schematic diagram of the role of miR-122 in slowing down SL neuron maturation. Data are represented as mean ± SEM. Biological replicates are distinguished by circles in the bar plots (A, C, E). One-way ANOVA (A, bottom left); Two-way ANOVA (A, bottom right; C; E); Fisher’s chi-square test (C). *p < 0.05, **p < 10^-2^, ***p < 10^-3^, ****p < 10^-4^.

## Discussion

Our results reveal that miR-137 and miR-122 control cortical development by acting on pre- and post-mitotic cells, respectively. MiR-137 acts on basal progenitors, maintaining their proliferative state, including through upregulation of the extracellular matrix receptor *Cd63* and repressing *Myt1l*, a transcription factor promoting neuronal identity. On the other hand, miR-122 slows down neuronal maturation pace.

We identify miR-137 and miR-122 as enriched in ferret oSVZ during SL neurogenesis. In the oSVZ, basal progenitors highly proliferate to sustain cortical expansion and folding (Pilz et al., 2013; Dehay et al., 2015). Recent studies showed that both BP types (namely bRG and IP) display an increased proliferative capacity (Kalebic et al., 2019), mainly through interactions with the extracellular matrix (Arai et al., 2011, Fietz et al., 2012). Here, we show that miR-137 maintains BPs in a proliferative state, not only through the maintenance of cell cycle genes and repression of neuronal genes, but also through the induction of an extracellular matrix receptor (*Cd63*) mediating the response to integrin signaling pathway, which is known to promote proliferation of BP (e.g. Integrin avb3 and b1 pathways; Fietz et al., 2010; Kalebic et al., 2019; Stenzel et al., 2014).

Progenitor division mode changes as corticogenesis proceeds. SL neuron generation mostly relies on BP, whereas a significant proportion of DL neurons are born directly from AP. Our data shows that miR-137 triggers BP proliferation both at the peak of BP-derived neurogenesis (i.e. during SL neuron generation) as well as at earlier time points (i.e. during DL neuron generation). This supports an evolutionary role for this miR in the disproportionate increase in SL neuron numbers in gyrencephalic compared to lissencephalic species.

The protracted duration of corticogenesis in more evolved species is reflected in both longer progenitor cell cycle duration (Bystron et al., 2006; Caviness et al., 1995; Rakic, 2009) and longer neurogenic period (Lewitus et al., 2014; Wilsch-Brauniger et al., 2016). In addition, postmitotic neuronal maturation is longer in more evolved species and human neurons transplanted into the mouse cortex, retaining a slower developmental pace (Linaro et al., 2019). In our study, we show that miR-122 acts postmitotically by modifying the transcriptional profile of newborn neurons during SL neurogenesis, along with a slower differentiation and migration pace and a shift in molecular identity from L4 to L2/3. In evolutionary terms, the lengthened maturation has been associated with a protracted differentiation of synapses and dendrites, increased synaptic plasticity, circuit integration and specialization, which could underlie the extended postnatal plasticity of neocortical circuits (Petanjek et al., 2011; Gould, 1992).

Non-coding RNAs have emerged over the last decade as a critical source of transcriptional control across species (Fernandez et al., 2016, Florio et al., 2017). The proportion of non-coding RNAs per genome size increases across evolution (Liu et al., 2013; Taft and Mattick, 2003; Taft et al., 2007) as do numbers of miRs and the length of their targeting region (Meunier et al., 2013). The regulatory importance of miRs is particularly appreciable during neocortical development. Several studies have shown that the absence of miRs during cortical development leads to severe malformations (Choi et al., 2008; Cuellar et al., 2008; Damiani et al., 2008; Davis et al., 2008; De Pietri Tonelli et al., 2008; Kim et al., 2007; Stark et al., 2008). Several conditional knock-down Dicer models used to study the role of miRs during neurogenesis of excitatory cortical neurons found that absence of miRs leads to premature neurogenesis, depletion of progenitors and migratory defects of newborn neurons. Moreover, gyrencephalic species show an enriched array of miRs (Jönsson et al. 2015, Moreau et al. 2013), in particular across the different germinal layers (Arcila et al., 2014), where they provide a major contribution to neuronal expansion. In our study we found that oSVZ enriched miR-137 and miR-122 modify the transcriptional landscape of the mouse basal progenitors and newborn neurons.

MiR-137 acts during SL neurogenesis, whereas miR-122 is involved in SL neuronal differentiation, triggering molecular dynamics happening in gyrencephalic species, which contribute pre and postmitotically to an evolutive expansion of the L2/3 identity. We think these findings unveil a parsimonious mechanism to orchestrate in time and space the neuronal expansion and differentiation, providing key insights into how non-coding RNAs transcriptionally shape developmental programs in the cortex evolution.

It would be interesting for future studies to investigate the mechanisms through which changes in the pace of molecular differentiation programs leads to distinct cellular fates.

## AKNOWLEGEMENTS

We thank Davide De Pietri for providing plasmids and Natalia Bauman for her contribution to the single-cell analyses. We are thankful to Audrey Benoit and Wafae Adouan and Greta Limoni for technical assistance and Didier Chollet and Mylene Docquier of the Genomics Platform of the University of Geneva. We thank Jabaudon and Dayer members of the laboratory for suggestions on the manuscript. The Jabaudon Laboratory is supported by the Swiss National Science Foundation and the Brain and Behavior foundation. E.K. is supported by a grant from the Machaon Foundation. J.P. is supported by the Public Instruction Department, Geneva. A.D., U.T., M.N., N.M., L.dV. are supported by the Swiss National Foundation Synapsy (grant 51NF40-158776), A.S is supported SNSF, scholarship SNF (175830), R.J.P. is supported, in part, by funds from the Botnar Research Centre for Child Health Multi-Investigator Project, the European Research Council (851021), the EMBO Young Investigator Program (4217), the Swiss National Science Foundation (31003A_175830), ETH domain Personalized Health and Related Technologies (PHRT-203), the National Centres of Competence – Molecular Systems Engineering, and the ETH Zurich (ETH-27 18-2) and V.B. is supported by European Research Council (309633), Spanish Research Agency (SAF2015-69168-R, PGC2018-102172-B-I00 and through the “Severo Ochoa” Program for Centers of Excellence in R&D, ref. SEV-2017-0723).

## AUTHOR CONTRIBUTIONS

A.D., V.B. and U.T. performed the experimental design; U.T. performed the experiments with help from E.K., M.N., and N.M., L.dV; Transcriptomic analyses were performed by U.T. and E.K. with the help of J.P.; U.T. and E.K. performed the data analysis and display; U.T and A.S. performed the cloning of expression vectors; Electrophysiological experiments were performed by N.M.; U.T. wrote the manuscript with the help of A.D., D.J, and the other authors. We dedicate this manuscript to the memory of Alexandre Dayer, who passed away before completion of this work.

## MATERIALS AND METHODS

### Experimental model and subject details

All experimental procedures were approved by the Geneva Cantonal Veterinary Authority and performed according to the Swiss law. Embryonic day (E) 0.5 was established as the day of vaginal plug. Wild-type CD1 mice were provided by Charles River Laboratories. Male and female embryos between E12.5 and E15.5 were used for the *in utero* electroporations, and pups between postnatal day (P) 0 and P21 for the postnatal experiments. Pregnant dams were kept in single cages and pups were kept with their mothers until P21, in the institutional animal facility under standard 12:12 h light / dark cycles.

### In *utero* electroporation

Timed pregnant CD1 mice were anaesthetized with isoflurane (5% induction, 2.5% during the surgery) and treated with the analgesic Temgesic (Reckitt Benckiser, Switzerland). Embryos were injected unilaterally with 700 nl of DNA plasmid solution (diluted in endofree PBS buffer and 0.002% Fast Green FCF (Sigma)) into the lateral ventricle. Embryos were then electroporated by holding each head between circular tweezer-electrodes (5 mm diameter, Sonidel Limited, UK) across the uterine wall, while 5 electric pulses (35 V for E12.5, 40 V for E13.5, 45 V for E14.5, 50 V for E15.5, 55 V for E17.5 and E18.5, 50 ms at 1 Hz) were delivered with a square-wave electroporator (Nepa Gene, Sonidel Limited, UK).

### Plasmids

Injected plasmids were: pUB6-GFP and pUB6-TOM (0,5 mg/ml); pSilencer-U6-scram, pSilencer-U6-miR-137 and pSilencer-U6-miR-122 (1 mg/ml); dUP-Cd63 and dUP-Myt1l (2 mg/ml, subcloned from double UP mClover to Scarlet, Addgene #125134, Taylor et al., BioRxiv 2019); DFRS, DFRS-137 and DFRS-122 (1 mg/ml, subcloned from DFRS, De Pietri Tonelli et al., 2006).

A vector backbone pSilencer 2.1 was used to clone the pSilencer-U6-miR-137 and the pSilencer-U6-miR-122. MiR-137 and miR-122 sequence is flanked by BamHI and HindIII excision sites, allowing the insertion into the pSilencer 2.1-U6 neo Vector (already linearized; Ambion) using the In-Fusion Kit (Clontech). The mir-137 and miR-122 sequence was thus under the control of the constitutive human RNA Polymerase III promoter U6.

The Dual-Fluorescent-GFP-Reporter/mRFP-Sensor plasmid (DFRS) is able to detect strong and weak expressions of miRNAs at a single cell level (De Pietri Tonelli et al., 2006). With The low endogenous level of miR-137 and miR-122 can be detected, as well as the effect of the gain of function, which allows the validation of the pSilencer-U6-miR-137 and the pSilencer-U6-miR-122 plasmids. In DFRS, both the red fluorescent protein (RFP) gene and the green fluorescentprotein (GFP) gene were driven by the identical simian vacuolating virus 40 (SV40) promoter, which induced a constitutive expression of the genes under this promoter (Oellig and Seliger, 1990). The GFP was the reporter, and thus was not affected by any miRNA. The 3’UTR of RFP gene contained the control cassette (unaffected by any miRNA), or the miR-137 or miR-122 target cassette (sensible to the expression of miR-137 or miR-122). The RFP transcript was thus the sensor, its translation being affected (DFRS137 or DFRS122) or not (DFRS control) by the presence of miR-137 or miR-122 in the cell. The DFRS control plasmid was constructed from the pGEM shuttle plasmid, which contained a control cassette (Unc-54 3’UTR, C. elegans), that did not include any miRNA active 3’UTR. Thus, this control cassette was not targeted by any miRNA from the mouse genome. This cassette was amplified (pGEM 3’UTR cassette primers) then removed from the pGEM shuttle plasmid using EcoRI and NotI enzymes and annealed into the DFRS empty plasmid pre-amplified (DFRS empty plasmid primers) and digested by the same enzymes (In-Fusion Kit; Clontech). The control cassette was thus placed on the 3’UTR of the RFP gene.

The DFRS137 and DFRS122 were the sensors of the level of miR-137 and miR-122 in the cell. For this, the 3’UTR of the RFP gene in the DFRS plasmid contained two perfect complementary sequence of miR-137 or miR-122 in tandem. Thus, these sequences were recognized and targeted by miR-137 or miR-122, which induced the degradation of the messenger RNA of the RFP. First, the primers for the complementary sequence of miR-137 or miR-122 were designed (miR-137 or miR-122 target cassette primers), flanked by PacI and FseI restriction sites. The cassette was then inserted by digestion and ligation into the pGEM shuttle plasmid. The complete cassette containing the 3’UTR information was amplified (pGEM 3’UTR cassette primers), digested by EcoRI and NotI and ligated to the DFRS empty plasmid, pre-amplified (DFRS empty plasmid primers) and pre-digested by the same enzymes (In Fusion Kit; Clontech). The miR-137 or miR-122 complementary sequences were thereby located on the 3’UTR of the RFP gene.

A vector backbone (pUG001) was constructed to allow for temporal control of gene expression (*CAG-loxP-mClover3-polyA-loxP-mScarlet-MCS-polyA*). Upon plasmid delivery, only mClover3 is constitutively expressed. As this gene is flanked by loxP sites, subsequent delivery of Cre protein results in the excision of mClover3 sequence, allowing for expression of mScarlet and the gene of interest (cloned at the MCS). A vector backbone (Double UP, Addgene #125134, Taylor et al., 2020) was used to allow for temporal control of gene expression (*CAG-loxP-mClover3-polyA-loxP-mScarlet-MCS-polyA*). Genes of interest were PCR amplified from a cDNA library from mouse embryonic brain RNA extracted at E14.5.

Double UP plasmid was modified to permit constitutive expression of the gene of interest, and its ablation upon Cre activation, by including a T2A site downstream of mClover3. The modified plasmid (pUG015) was digested with BglII (ThermoFisher, FD0083) and Gibson Assembly (NEB, E2611S) was used to introduce Cd63 (pUG016) and Myt1L (pUG018) downstream of T2A site.

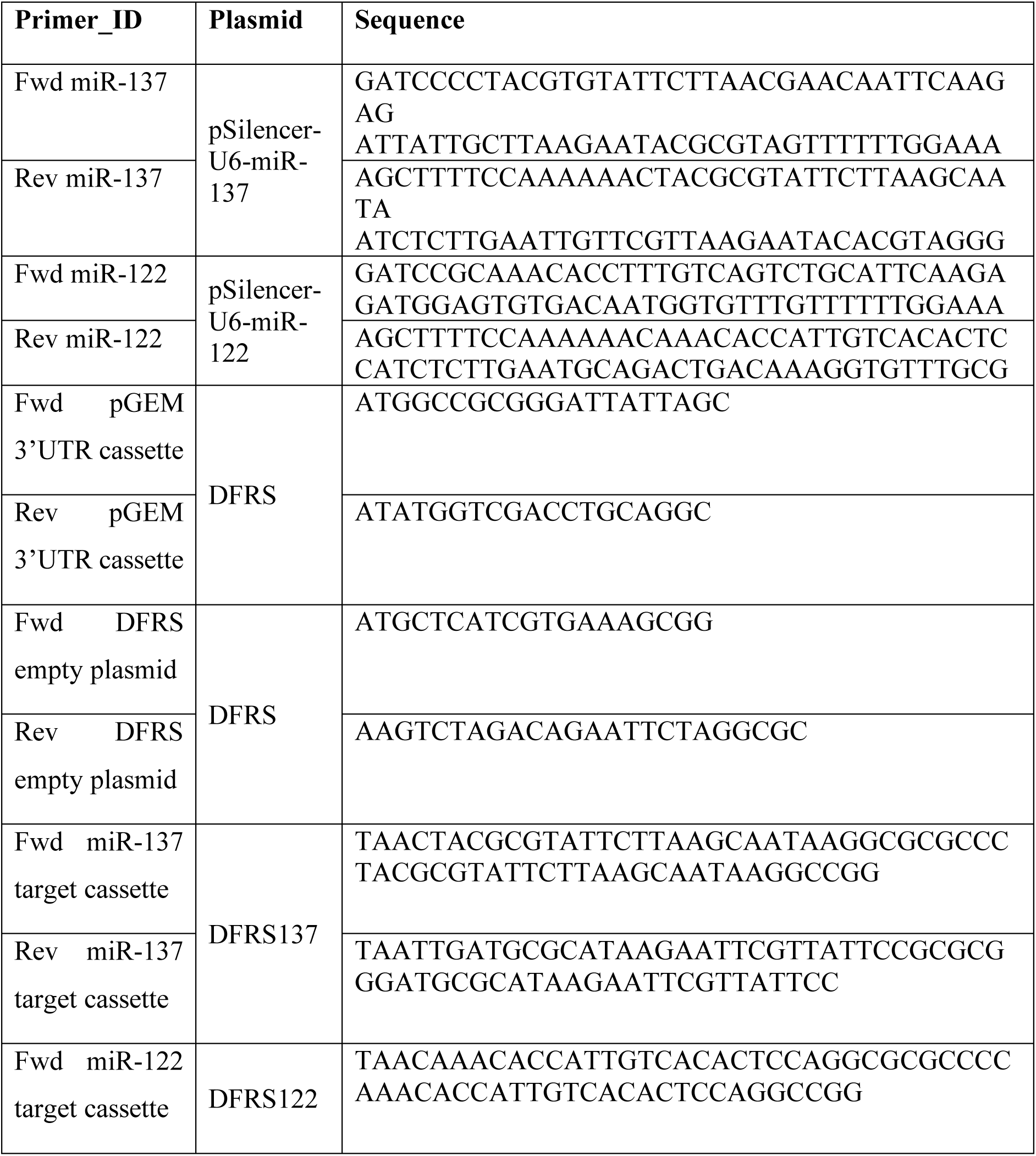
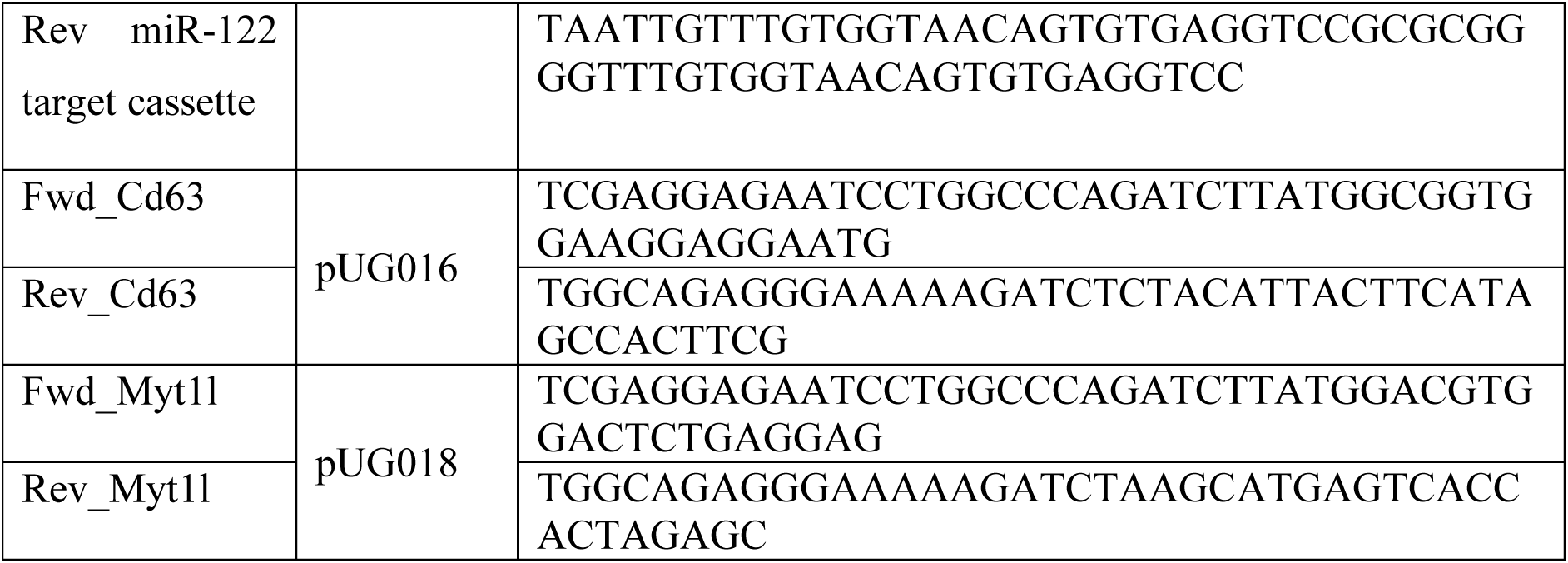

### Injections and continuous drugs administration (EdU)

For chronic administration, an osmotic pump (Alzet, #1003D) was filled with 10 mg/ml solution of EdU (Sigma) was prepared in 1:1, DMSO : water and placed in the peritoneal cavity at the end of the surgery or at given gestational day (Telley et al., 2016). For single-pulse labeling, a single dose of 10 mg/kg of animal weight of EdU (10 mg/ml in water) was administered intra-peritoneally.

### Retrograde labeling

Anesthetized pups were placed in a stereotaxic apparatus at postnatal day (P) 9 and injected with red Retrobeads (Rbeads) IX from Lumafluor in the primary somatosensory cortex (S1) (200 nl; coordinates from the lambda: anteroposterior: 3 mm, mediolateral: 3 mm).

### Immunohistochemistry and imaging

Embryos were collected 24, 48 and 72 hrs following *in utero* electroporation and post-fixed overnight in 4% paraformaldehyde (PFA, Sigma) at 4°C. Postnatal mice from P0 were perfused intracardially with 4% PFA and post-fixed overnight in 4% PFA at 4°C. 80 μm coronal sections were performed using a vibratome (Leica, #VT100S). Sections were permeabilized in PBST (0.3% Triton X-100, diluted in PBS 1X) and incubated for two hrs at room temperature in blocking solution (10% Horse Serum Albumine in PBST), then overnight at 4°C with primary antibodies. Treatment with HCl 2N at 37°C for 30’ was performed before incubation with standard blocking solution for KI67 immunohistochemistry. Treatment with Na citrate pH 6 at 80°C for 40’ was performed before incubation with standard blocking solution for RORB immunohistochemistry.

Sections were rinsed three times in PBST and incubated for 2 hrs at room temperature with corresponding secondary antibodies (1:500, Life Technologies). Three washes in PBST were performed, followed by 10 min incubation with Hoechst staining solution (1:5000 in PBS 1X, Life Technologies) to label nuclei, before dry mounting on slides with Fluoromount (Sigma). For imaging, the putative primary somatosensory cortex (S1) was used as region of study for all the experiments. Images were acquired on Eclipse 90i epifluorescence microscope (Nikon) or on LSM 700 confocal laser scanning microscope (Carl Zeiss).

### Antibodies

Chicken anti-GFP (1:2000; Abcam, #AB13970); Goat anti-TOM (1:300; Sicgen, #AB8181-200); rabbit anti-RFP (1:100; Abcam, #AB62341); Mouse anti-PAX6 (1:300; Thermoscientific, #MA1-109); Rat anti-EOMES (1:500; Invitrogen, #14-4875-82); Rat anti-EOMES (1:300; eBioscience, #144875-82); Rabbit anti-KI67 (1:250; Abcam, #AB15580); Rabbit anti-NEUROD2 (1:1000; Abcam, #AB104430); Mouse anti-RORB (1:200; Perseus Proteomics, #PP-N7927-00); Rabbit anti-CUX1 (1:250; Santa Cruz, #sc13024), Rabbit anti-CASPASE3 (1:3000; Cell signaling technology, #9662S). All secondary antibodies were 488/555/647 conjugated (1:500, Invitrogen).

### Time-lapse imaging

Acute slices were prepared at E17.5 from brains electroporated into the dorsal pallium at E14.5 with either *pSilencer scram*, *pSilencer miR-137* or *pSilencer miR-122* along with *pUb6-tdTOM* (reporter) plasmids. Briefly, brains were dissected out in ice-cold HBSS (Gibco), embedded in 3% LM-agarose (Roth) and cut 250 µm-thick with a vibratome (Leica VT1000S). Slices were then transferred on Millicell inserts (Merck Millipore) on NBM medium in incubator at 37°C for at least 2h recovery before being placed in a Fluorodish (WPI) for imaging. In all the experiments, scramble and miR-electroporated cell were imaged simultaneously. Statistical significance was set at α<0.05. All calculations were done using Excel (version 16.31) and statistics were done with GraphPad Prism software (version 8.1.2). Normality of the samples was assessed with D’Agostino-Pearson test and non-parametric tests used when criteria were not fulfilled.

#### Multipolar-bipolar transition and radial migration imaging

Imaging was achieved with a live confocal microscope (Nikon A1r) equipped with long-working distance 20x or 40x objectives (0.45 and 0.6, respectively, CFI ELWD Plan Fluor, Nikon). The microscope chamber was kept at 37°C, with 25l/h continuous flux of 5% CO_2_ humidified at 95%. 50µm-thick z-stacks (3µm-stepped) were acquired with resonant laser scanning every 10min for 12h. In all the experiments, scramble and miR-electroporated cells were imaged simultaneously. At the end of the imaging session, stacks were piled into maximal intensity projections and time sequences corrected for drift with ImageJ (StackReg). To measure the multipolar to bipolar switch (**Fig. 4F**), a box of 300×x150µm was drawn at the transition zone with the upper border aligned on the IZ/deep layers boundary. All multipolar-shaped cells contained in this box were followed along 12 h and the percentage of cells transiting from multipolar morphology to bipolar shape was calculated. Cell movements were manually tracked and dynamic data extracted with ImageJ (MTrackJ). A cell was considered pausing when its movement along 10 min was under 12µm. Average speed is the distance achieved by a cell divided by the time taken to travel it, movement speed is the same calculation but not considering the events of pausing and directionality is the distance in straight line between the start and end point of a cell divided by the distance it travelled between these two points (**Fig.4F** and **S4B**). After manual tracking and data extraction, a random selection of an equal number of cells per slices was applied to avoid overrepresentation bias. All calculations were done using Excel (version 16.31) and statistics were done with GraphPad Prism software (version 8.1.2), with statistical significance set at α<0.05. Normality of the samples was assessed with D’Agostino-Pearson test and non-parametric tests used when criteria were not fulfilled.

### Tissue microdissection, cell sorting and RNA sequencing

#### Single-cell RNA sequencing

##### Embryonic cortex

Tissue collection was performed as in Telley et al., 2019. Briefly, pregnant females were sacrificed, and embryos at E15.5 (*n* = 6 Scramble and *n* = 6 miR-137 embryos) and E17.5 (*n* = 6 Scramble and *n* = 6 miR-122 embryos) after E14.5 *in utero* electroporation extracted in ice-cold HBSS. 300µm-thick acute coronal brain sections were cut after embedding in 4% low melt-agar using a vibratome (Leica, #VT1000S) under RNase-free conditions. The putative S1 was microdissected using a Dissecting Scope (Leica, #M165FC) and incubated in 0.05% trypsin at 37°C for 5 minutes. Following tissue digestion, cells were incubated in fetal bovine serum and manually dissociated via gentle pipetting. Cells were then centrifuged for 5 minutes at 500 rpm, resuspended in 1 mL of HBSS, filtered using a 70 μm-pored cell strainer (ClearLine, #141379C) and incubated for 10 minutes at 37°C with Hoechst (0.1 mg/mL).

##### Postnatal cortex

Tissue collection was performed as in Klingler et al., Curr Biol 2019. Briefly, 300 μm acute coronal brain sections were cut on a vibratome and S1 was microdissected as described above in ice-cold oxygenated artificial cerebrospinal fluid (ACSF) under RNase-free conditions.

P3: *n* = 4 E14 GFP electroporated pups; *n* = 4 E15.5 GFP electroporated pups. P7: *n* = 4 E14 GFP electroporated pups; *n* = 4 E15.5 GFP electroporated pups; *n* = 4 E14.5 Scramble + GFP electroporated pups; *n* = 4 E14.5 miR-122 + GFP electroporated pups; *n* = 4 E14.5 miR-137 + GFP electroporated pups

For L4 *vs*. L2/3 comparison at P3 and P7 (i.e. E14 *vs*. E15.5 GFP electroporated pups), layers were microdissected following the same procedure as described above with further separation of superficial (SL) and deep (DL) layers. 5000 to 10’000 SL and DL cells were next dissociated by incubating microdissected samples in 0.5 mg/mL Pronase (Sigma, #P5147) at 37°C for 10 minutes, followed by a 3 minute inactivation in 5% bovine serum albumin, washes in ACSF and manual trituration using pulled glass pipettes of decreasing diameters. Cells were then centrifuged for 10 minutes at 500 rpm, resuspended and filtrated using a 70 μm-cell strainer (ClearLine, #141379C) and incubated for 10 minutes at 37°C with Hoechst (0.1 mg/mL).

Singlet GFP^+^/Hoechst^+^ embryonic and postnatal cells were sorted using a Beckman Coulter Moflo Astrios FAC-sorter according to their Forward and Slide scattering properties, and their negativity for Draq7^TM^ (Viability dye, far red DNA intercalating agent, Beckman Coulter, #B25595). 5000 to 10’000 cells were FAC-sorted for each experiment. 3 µl of C1 Suspension Reagent (Fluidigm) was added to 10 μl of FACsorted cells, which were captured into 800 well-AutoPrep integrated fluidic circuit (IFC) designed for 10 to 17 μm diameter-cells (Fluidigm HT800, #101-4982) for embryonic cells, and for 10 to 17 μm diameter-cells (Fluidigm HT800, #100-57-80) for postnatal cells, and imaged using the ImageXpress Micro Widefield High Content Screening System (Molecular Devices®). cDNA synthesis and preamplification was processed following the manufacturer’s instructions (C1 system, Fluidigm). cDNA libraries were prepared using Nextera XT DNA library prep kit (Illumina), quality control was done using 2001 Bioanalyzer from Agilent and sequenced using HiSeq 2500 sequencer for E14.5 electroporated pups, and HiSeq 5000 sequencer for E14 and E15.5 electroporated pups.

#### Bulk RNA sequencing

Brains were collected from E14.5 *in utero* electroporated miR-122 pups at P7 (*n* = 3 pups). S1 was microdissected following the same procedure as described above with further separation of superficial (SL) and deep (DL) layers. 5000 to 10’000 SL and DL cells were dissociated and FAC-sorted as for single-cell RNA sequencing. RNA from each sample was extracted using Total RNA Isolation System kit (Promega SV) and quality control was done using 2001 Bioanalyzer from Agilent. cDNA libraries were obtained using SMARTseq v4 kit (Clontech, # 634888) and sequenced using HiSeq 2500 sequencer. All single cell RNA capture, library preparations and sequencing experiments were performed at the Genomics Core Facility of the University of Geneva.

### Electrophysiology

In *utero* electroplated (at E14.5) CD1 mice from P20-P26 were used for electrophysiological recordings. Mice were anesthetized with isoflurane and decapitated to dissect out the brains and were immediately transferred into ice-cold sucrose cutting solution equilibrated with 95% O_2_ and 5% CO_2_ containing (in mM) Sucrose (75), NaCl (85), CaCl_2_ (0.5), MgCl_2_ (4), NaHCO_3_ (24), KCl (2.5), NaH_2_PO_4_ (1.25) and glucose (25). Three hundred µm thick coronal slices were cut using a vibratome (Leica VT 1200S).

*For another set of experiments, embryonic slices were obtained at E17.5 from electroporated embryos (E14.5). Briefly, embryos were surgically dissected and transferred in ice-cold HBSS (Gibco, 14175-053), embedded in 2% agarose solution, and four hundred µm thick coronal slices were cut in HBSS using a vibratome (Leica VT 1200S)*.

Slices were incubated at 35 °C for 20 min in a slice recovery chamber filled with artificial cerebrospinal fluid (ACSF) containing (in mM) NaCl (125), CaCl_2_ (2.5), MgCl_2_ (1), NaHCO_3_ (26), KCl (2.5), NaH_2_PO_4_ (1.25) and glucose (25). Slices were kept at room temperature in a recovery chamber until recording. For recording, slices were transferred to a recording chamber continuously perfused with oxygenated ACSF that was maintained at 30±0.1°C *(34±0.1°C, for embryonic slices)* using an in-line heating system (TC-01, Multichannel systems). All the recordings were carried out into the somatosensory cortex; the barrels in L4 were used as a visual landmark for identification. *Embryonic recordings were carried out in intermediate zone.* Cortical layers 2/3 and 4 neurons were visualized by using an upright microscope and camera system (BX51WIF, Olympus, and SciCam Pro CCD camera, Scientifica), equipped with a 40x water-immersion objective, infrared/differential interference contrast (DIC) optics, and epifluorescence (GFP filter set and a LED source: COO-LED2LLG-470-565, CoolLED). Electroporated neurons were identified using GFP expression and were used for further recordings. Whole-cell recordings were obtained with recording pipettes of resistance between 3-5 MΩ. The recording pipettes were pulled using borosilicate glass capillaries (1.5 mm OD, GC150TF-7.5, Harvard Instruments) on Zeitz DMZ puller (Zeitz instrument). Pipettes were filled with an internal solution containing (in mM) CH_3_KO_3_S (140), MgCl_2_ (2), NaCl (4), creatine phosphate (5), Na_2_ATP (3), GTP (0.33), EGTA (0.2) and HEPES (10) adjusted to 295 mOsm and pH 7.3 with KOH.

Whole-cell voltage-clamp recordings were obtained using a Multiclamp 700B amplifier (Axon Instruments) filtered at 3 kHz and digitized at 20 kHz (NI-6341, National Instruments Board and Igor, WaveMetrics). Neurons were held at -70 mV in voltage-clamp mode and a pulse of -4 mV was given at 0.1 Hz to monitor series resistance (R_s_). Neurons with a stable R_s_ and stable resting membrane potential (RMP) negative than -60 mV were subjected to a battery of current injection protocol to study electrophysiological properties. I_h_ currents were recorded in voltage-clamp using a - 40 mV step (500 ms) and were calculated by the difference of current between the beginning and the end of the voltage step. Spontaneous excitatory postsynaptic currents (sEPSCs) were recorded at V_h_=-70 mV 1 µM of SR95531 hydrobromide (Tocris, Cat no. 1262) was bath applied to block inhibitory neurotransmission. Firing characteristics were studied in the current-clamp mode by injecting incremental steps of depolarizing currents from+50 pA to 500 pA for 500 ms.

### Quantification and statistical analysis

#### Histological analyses

Zen (Zeiss) and ImageJ softwares were used to analyze images. All results are shown as mean ± SEM, except when indicated otherwise. For statistical analyses, the following convention was used: *: p < 0.05, **: p < 0.01, ***: p < 0.001. ‘‘Student’s t-test’’ refers to the unpaired test. For embryonic analyses, differently electroporated embryos from the same mother were used to reduce plug timing variability. Experiments were cross-quantified blindly (i.e., the investigator was unaware to which of the experimental conditions the sections were belonging). Where indicated, scram, miR-137 and miR-122 were analyzed together in order to perform more robust statistical analysis (One-way Anova in place of Unpaired t-Test).

Figures 1B-D, 2D, S2G: Three sections for each brain electroporated with pUB6-TOM + pSilencer-U6-scram (number of brains, *n* = 3) or pUB6-TOM + pSilencer-U6-miR-137 (number of brains, *n* = 3) or pUB6-TOM + pSilencer-U6-miR-122 (number of brains, *n* = 3) at E12.5 or E14.5, were used to quantify the number of KI67^+^, NEUROD2^+^, PAX6^+^ and EOMES^+^, cells among the fraction of TOM^+^ cells 36, 48 or 72hr after IUE. Three sections for each brain electroporated with pSilencer-U6-scram + dUP-Cd63 (number of brains, *n* = 3) or pSilencer-U6-scram + dUP-Myt1l (number of brains, *n* = 3) at E14.5 were used to quantify the number of KI67^+^, NEUROD2^+^, PAX6^+^, EOMES^+^, cells among the fraction of GFP^+^ cells 48hr after IUE.

**Figure 1B**: percentage of VZ+SVZ KI67^+^ in E16.5 scram: 47.91 ± 1.6, E16.5 miR-137: 75.11 ± 3.12, E16.5 miR-122: 50.37 ± 2.31. Percentage of KI67^+^ in VZ E16.5 scram: 56.81 ± 1.67, VZ E16.5 miR-137: 34.69 ± 2.3, VZ E16.5 miR-122: 53.68 ± 1.17, SVZ E16.5 scram: 43.18 ± 1.67, SVZ E16.5 miR-137: 65.3 ± 2.3, SVZ E16.5 miR-122: 46.31 ± 1.17. A one way-ANOVA with Dunnett’s post hoc test was used when required.

**Figure 1C**: percentage of NEUROD2^+^ in E17.5 scram: 49.44 ± 1.24, E17.5 miR-137: 63.09 ± 1.4, E17.5 miR-122: 52.9 ± 0.67. A one way-ANOVA with Dunnett’s post hoc test was used when required.

**Figure 1D**: ratio of PAX6^+^/EOMES^+^ in E16.5 scram: 1.03 ± 0.02, E16.5 miR-137: 3.76 ± 0.18, E16.5 miR-122 (not shown): 1.25 ± 0.07, E14 scram: 1.13 ± 0.18, E14 miR-137: 3.25 ± 0.33. A one way-ANOVA with Dunnett’s post hoc test was used for E16.5. A student-T test was used for E14.

**Figure 2D**: percentage of KI67^+^ in E16.5 scram: 47.91 ± 1.6, E16.5 miR-137: 75.11 ± 3.12, scram+Myt1l-gfp: 34.85 ± 1.02, scram+Cd63-gfp: 73.31 ± 1.79. Percentage of NEUROD2^+^ in E16.5 scram: 49.46 ± 1.17, E16.5 miR-137: 35.14 ± 1.75, scram+Myt1l-gfp: 71.5 ± 2.26, scram+Cd63-gfp: 36.31 ± 1.52, E16.5 miR-137: 3.76 ± 0.18. Ratio of PAX6^+^/EOMES^+^ in E16.5 scram: 1.03 ± 0.02, E16.5 miR-137: 3.76 ± 0.18, scram+Myt1l-gfp: 1.1 ± 0.1, scram+Cd63-gfp: 2.05 ± 0.07, E16.5 miR-137: 3.76 ± 0.18. The same scram, miR-137 and miR-122 brains were used for the Fig. 1B or 1C or 1D and Fig. 2D. Scram, miR-137, miR-122, scram+Myt1l and scram+Cd63 were analyzed together to minimize biological variability. A one way-ANOVA with Dunnett’s post hoc test was used when required.

**Figure S1D**: three to four sections for each brain electroporated at E14.5 with DFRS or DFRS137 or DFRS122 and pSilencer-U6-miR-122 (number of brains, *n* = 3) or pSilencer-U6-miR-137 (number of brains, *n* = 3) were used to quantify the ratio of GFP^+^/RFP^+^ cells at E15.5. The analyses have been carried blindly. Ratio of GFP^+^/RFP^+^ in DFRS: 0.97 ± 0.08, DFRS137: 0.71 ± 0.03, DFRS137+pSil-miR-137: 0.25 ± 0.03, DFRS122: 0.42 ± 0.02, DFRS122+pSil-miR-122: 0.25 ± 0.002. A one way-ANOVA with Tukey’s post hoc test was used when required.

**Figures 3A**, **5A**: two to three sections for each electroporated brain were used to define the laminar position (Y coordinate) of electroporated cells at P7. Analyses were carried blindly. The Y value was normalized as percentage of distance from the WM. The cortex was divided in 10 bins and the mean value of frequency distribution was plotted in a bar graph. Dark-colored bins represent significant difference for the considered conditions. The Y values were plotted grouped by brain and the mean value and standard deviation were represented. Density plot and cumulative distribution were used to additionally display the Y values.

**Figure 3A**: IUE at E14.5 with pUB6-TOM and pSilencer-U6-scram (number of brains, *n* = 4) or pSilencer-U6-miR-137 (number of brains, *n* = 4). Mean value for Y position in E14.5 control: 36.13 ± 0.29, miR-137: 34.36 ± 0.27.

**Figure 5A**: IUE at E14.5 with pUB6-TOM and pSilencer-U6-scram (number of brains, *n* = 4) or pSilencer-U6-miR-122 (number of brains, *n* = 4). Mean value for Y position in E14.5 control: 36.13 ± 0.29, miR-122: 66.67 ± 0.79. The same scram brains were used for the Fig. 3A and Fig. 5A. Scram, miR-137 and miR-122 were analyzed together to minimize biological variability. A two-way ANOVA with Bonferroni’s post-hoc test was used for bin analysis; a one-way ANOVA was used for mean Y position.

**Figure 3B**: three sections for each brain electroporated at E14.5 with pUB6-GFP and pSilencer-U6-scram (number of brains, n = 4) or pSilencer-U6-miR-137 (number of brains, n = 4) and chronically-delivered EdU at E16.5, were used to quantify the number of EdU cells among the total amount of GFP^+^ cells in L2/3 at P7. Percentage of EdU/GFP^+^ scram: 10.74 ± 0.74, miR-137: 27.55 ± 2.09. A student-T test was used.

**Figures 3D**, **5C**, **S3B**: three sections for each brain electroporated at E14.5 with pUB6-GFP and pSilencer-U6-scram (number of brains, *n* = 4) or pSilencer-U6-miR-137 (number of brains, *n* = 4) or pSilencer-U6-miR-122 (number of brains, n = 4) were used to quantify the number of RORB^+^ cells among the fraction of electroporated cells in L2/3 and L4 at P7. Ratio of RORB^+^ in E14.5 L2/3 control: 0.06 ± 0.01, L2/3 miR-137: 0.07 ± 0.008, L2/3 miR-122: 0.03 ± 0.004, L4 control: 7.29 ± 0.46, L4 miR-137: 2.53 ± 0.23, L4 miR-122: 0.62 ± 0.11. A two-way ANOVA with Bonferroni’s post-hoc test was used.

**Figures 3E**: coronal sections from at least 3 different brains electroporated at E14.5 with pUB6-GFP and pSilencer-U6-scram (total of 676 cells) or pSilencer-U6-miR-137 (total of 969 cells) were used to quantify the percentage of electroporated cells displaying an apical dendrite at P7 in L4. Percentage of cells with apical dendrite in E14.5 scram: 9.99 ± 1.33, E14.5 miR-137: 16.69 ± 1.58. A student-T test was used.

**Figure 3G**: three sections for each brain electroporated at E14.5 with pUB6-TOM and pSilencer-U6-scram (number of brains, *n* = 4) or pSilencer-U6-miR-137 (number of brains, *n* = 4) and injected with retrobeads (Lumafluor) at P9, were used to quantify the number of retrobeads^+^ cells among the fraction of TOM^+^ cells at P11. Ratio of retrobeads^+^ in L2/3 scram: 1 ± 0.14, L4 scram: 0.03 ± 0.01, L4 miR-137: 0.23 ± 0.09. A two-way ANOVA with Tukey’s post-hoc test was used when required.

**Figures 4E, S4C**: three sections for each brain electroporated at E14.5 with pUB6-TOM or pUB6-GFP and pSilencer-U6-scram (number of brains, *n* = 3) or pSilencer-U6-miR-122 (number of brains, *n* = 3) or pSilencer-U6-miR-137 (number of brains, *n* = 3, not shown) were used to quantify the number of TOM^+^ or GFP^+^/SATB2^+^ cells in each cortical layer at E17.5. The analyses have been carried blindly.

**Figure 4E**: Distribution of scram cells: (VZ/SVZ: 15.05 ± 0.47, IZ: 50.92 ± 0.7, L5/6: 21.5 ± 0.25, SL: 12.51 ± 1.03), miR-122 (VZ/SVZ: 35.91 ± 1.71, IZ: 61.07 ± 1.48, L5/6: 1.86 ± 0.04, SL: 1.14 ± 0.25), miR-137 (VZ/SVZ: 8.67 ± 1.06, IZ: 44.21 ± 1.23, L5/6: 17.66 ± 0.84, SL: 29.45 ± 1.9). A two-way ANOVA with Sidak’s post hoc test was used.

**Figure S4C**: SATB2 intensity of scram cells: (Global: 0.48 ± 0.04, SVZ: 0.11 ± 0.02, IZ: 0.47 ± 0.04, L5/6: 0.59 ± 0.05, SL: 0.61 ± 0.06), miR-122 (Global: 0.28 ± 0.02, SVZ: 0.09 ± 0.02, IZ: 0.29 ± 0.02, L5/6: 0.33 ± 0.02, SL: 0.45± 0.02). A student-T test was used for the global analysis. A two-way ANOVA with Sidak’s post hoc test was used for the layer-wise analysis. The fluorescence intensity (0-255 scale of 8-bit images) for SATB2^+^ of the counted cells has been plotted to show the expression of the marker.

**Figure 5E**: three sections for each brain electroporated at E14.5 with pUB6-GFP and pSilencer-U6-scram (number of brains, *n* = 3) or pSilencer-U6-miR-122 (number of brains, *n* = 3) were used to quantify the number of GFP^+^ cells in each cortical layer at P21. Analyses were carried blindly. Distribution of scram cells: (L2/3: 40.63 ± 2.85, L4: 57.95 ± 2.34, L5/6: 1.42 ± 0.52, WM: 0), miR-122 (L2/3: 55.81 ± 0.25, L4: 37.74 ± 0.74, L5/6: 6.14 ± 0.69, WM: 0.29 ± 0.29). A two-way ANOVA with Sidak’s post hoc test was used.

**Figure S5A**: three sections for each brain electroporated at E14.5 with pUB6-TOM with pSilencer-U6-scram (number of brains, *n* = 3) or pUB6-GFP with pSilencer-U6-miR-122 (number of brains, *n* = 3) were used to quantify the number of TOM^+^/EdU^+^ or GFP^+^/EdU^+^ cells in each SL or DL+WM at P7. Analyses were carried blindly. Distribution of scram cells: (SL: 99.94 ± 0.05, DL+WM: 0.05 ± 0.05), miR-122 (SL: 86.62 ± 0.63, DL+WM: 13.37 ± 0.63). A two-way ANOVA with Sidak’s post hoc test was used.

**Figure S5E**: three sections for each brain electroporated at E14.5 with pUB6-TOM and pSilencer-U6-scram (number of brains, *n* = 3) or pUB6-TOM/pUB6-GFP and pSilencer-U6-miR-122 (number of brains, *n* = 3) were used to quantify the number of CASP3^+^ cells at P7. Analyses were carried blindly. Percentage of scram cells: (1.67 ± 0.98), miR-122 (0.51 ± 0.51). A Unpaired t-Test was used.

### Single-cell transcriptomic analyses

Reads were mapped on mouse genome GRCm38 following the same pipeline described in Telley et al., Science 2019. Read 1, which contains the UMI sequence, was appended at the end of read 2 header; reads 2 were further mapped to the mouse genome with Tophat v2.0.13. Resulting alignment files in BAM format were processed with umi_tools (Smith, T., Heger, A. & Sudbery, I. UMI-tools: modeling sequencing errors in Unique Molecular Identifiers to improve quantification accuracy. Genome Res. 27, 491–499 (2017)) to deduplicate reads with identical UMI. Gene expression quantification was performed with R using summarizeOverlAP method of package GenomicAlignments. Only reads falling into exonic part of a gene are quantified (including 5’ and 3’ UTRs).

For single-cell RNA sequencing, each transcriptome was further associated to a manual brightfield picture annotation, where the presence of a single cell in the wells of the Fluidigm HT800 chips was checked. Only wells where a single GFP^+^ cell was observed were kept for further analyses (wells with no cell, cell(s) with convoluted shapes, multiple cells, or cell(s) with debris were excluded).

All bioinformatics transcriptomics analyses were performed using R programming language and Bioconductor packages as described below on reads per million (RPM) normalized (log_10_) gene expressions.

#### Single cell filtering

##### Embryonic single cells

Cells expressing less than 2000 genes, or more than 15% mitochondrial genes were excluded from the analysis (resulting in E15.5: *n* = 192 scramble cells, *n* = 222 miR-137 cells; E17.5: *n* = 131 scramble cells, *n* = 204 miR-122 cells).

##### Postnatal single cells

Cells expressing less than 1000 genes, or more than 15% mitochondrial genes were excluded from the analysis (resulting in E14.5 *in utero* electroporation, P7: *n* = 341 P7 scramble cells, *n* = 352 miR-137 cells, *n* = 269 miR-122 cells; E14 *in utero* electroporation: P3: *n* = 207 scramble cells, P7: *n* = 162 scramble cells; E15.5 *in utero* electroporation, P3: *n* = 211 scramble cells, P7: *n* = 229 scramble cells).

##### Clustering analysis of embryonic single cells

Seurat bioinformatics pipeline (https://cran.r-project.org/web/packages/Seurat/citation.html) was used to determine the most 2000 variable genes using the FindVariableFeatures function (selection method = vst) and to perform t-Distributed Stochastic Neighbor Embedding dimensional reduction (tSNE) from the top 10 principal components. The FindClusters function (resolution = 0.5) next revealed 4 clusters, which were assigned to apical progenitor (AP), basal progenitor (BP), immature neuron (N_0_), and mature neuron (N_1_) identities based on their gene expression analyzed using the FindAllMarkers function. Expression of cell type-specific transcripts described in Telley et al., Science 2019 was used to further validate the identified clusters (**Fig. 2A**, **Fig. 4A**).

*Pseudotime analysis of embryonic single cells* was performed for E17.5 scramble *vs*. miR-122 (**Fig. 4D**) experiments using a regularized ordinal regression method (*bmrm* R package) to predict differentiation status of each cell. The linear models were built using the scramble cells, ranking the genes according to their ability to predict each cell category (AP, BP, N_0_ and N_1_; for E17.5 experiment, AP and BP were pooled together as they represent a very small cluster), and re-optimized on the best 100 genes (top 50 and bottom 50), before cross-validations using leave-one-out method. Pseudo-differentiation prediction scores were then calculated for miR or human embryonic neurons from Nowakowski et al., 2017 (**Fig. 4H** and **Fig. S4D;** annotations from the original publication were used to define immature and mature neurons) cells by the linear combination of the top gene expression and compared to the cross-validation values of scramble cells.

##### Predictions of SL vs. DL miR-122 identity (Fig. S5B), L2/3 vs. L4 identity (Figs. 3C, 5B, S3A) or P3 vs. P7 L2/3 (or L4) identity (Fig. S5C)

A logistic regression model with regularization was used to build binary prediction models of: (1) microdissected SL *vs*. DL miR-122 cells (bulk RNA sequencing), (2) L2/3 (E15.5 *in utero* electroporated) *vs.* L4 (E14 *in utero* electroporated) P7 scram cells (single cell RNA sequencing), or (3) P3 *vs.* P7 L2/3 or L4 scram cells (single cell RNA sequencing). This implementation was provided by the hingeLoss function of the *bmrm* R package. This allowed to rank the genes based on their ability to predict the identity, and to re-train a new model on the best 100 genes (top- and bottom-50 genes). All model performances were addressed by leave-one-out (1) or 20-fold (2, 3) cross-validations on the subset of genes, which gave a prediction value to build receiver operating characteristic (ROC) curves. The layer / age prediction scores were then calculated for E14.5 *in utero* electroporated P7 miR *vs*. scram cells by the linear combination of the top gene expression. **Fig. 3C** and **5B**: percentage of scram cells: (L2/3: 53.07, L4: 46.92), miR-122SL (L2/3: 80.37, L4: 19.62), miR-122DL (L2/3: 66.66, L4: 33.33), miR-137 (L2/3: 66.19, L4: 33.8). A Fisher’s chi-square test was used.

*Differential gene expression analyses* between scram and miR-137 (E15.5: **Fig. 2**, P7: **Fig. 3**) or scram and miR-122 (E17.5: **Fig. 4**, P7: **Fig. 5**) cells were performed using logistic regression models for each cell type (hingeLoss function; *bmrm* R package). This allowed to rank the genes based on their ability to predict scram *vs*. miR identities, and to calculate an identity score based on the best 100 genes by 20-fold cross-validations. The areas under the ROC curve (AUC) were calculated based on the sensitivity / specificity of each model using the cross-validation values (roc.stat function).

All gene ontologies were performed using GSEA (Subramanian, A. *et al*. Gene set enrichment analysis: A knowledge-based approach for interpreting genome-wide expression profiles. *Proceedings of the National Academy of Sciences* 102, 15545–15550 (2005)).

### Electrophysiology data analysis

Neurons with R_s_ fluctuation of > 20% during recording were excluded from the analysis. Offline analysis of electrophysiological data was carried out using Igor pro (Wave metric), several electrophysiological parameters were manually computed as illustrated in **Fig. S5D**. The first AP elicited in response to threshold depolarizing current injection was used to calculate the single AP parameters. AP train elicited in response to the current injection of +500 pA for 1000 ms was used for calculation of spike frequency and other AP train parameters. For calculation of membrane properties, at least 12 consecutive sweeps were digitally averaged. The membrane time constant (tau) was computed by monoexponential fit to the first 100 ms after the current injection of -40 pA. The sag ratio was calculated using this equation: (V_min_ – V_end_)/V_min_; where V_min_ is the minimum voltage reached during the hyperpolarizing pulse of - 200 pA, and V_end_ is the final voltage reached at end of current injection. sEPSCs were detected and analyzed using Mini Analysis Program (Synaptosoft) script integrated into Igor pro. Prism 8 (GraphPad) was used for statistical analysis and preparation of the graphs. All the data are presented as mean±SEM.

**Figure 3F**: Coronal sections from at least 3 different brains electroporated at E14.5 with pUB6-GFP and pSilencer-U6-scram L2/3 (total of 13 cells) or L4 (total of 19 cells) or pSilencer-U6-miR-137 (total of 13 cells) were used to quantify the I_h_ current. Mean of cells with I_h_ in L2/3 scram: 59.85 ± 11.2, L4 scram: 27.32 ± 4.69, L4 miR-137: 43.53 ± 6.25. A student-t test was used.

---

**Figure S1:**
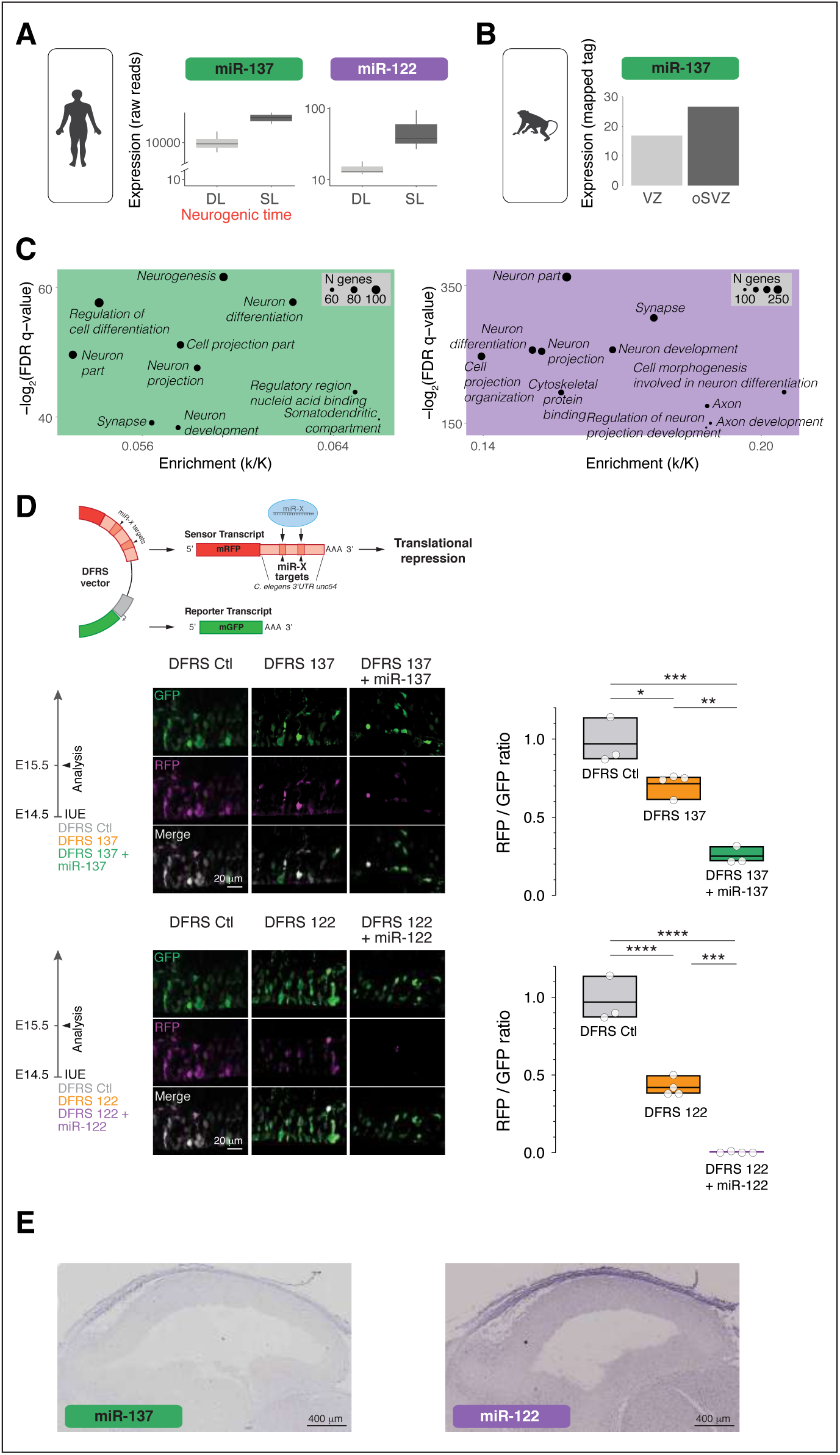
MicroRNAs 137 and 122 expression across species. (**A**) MiR-137 and miR-122 expression during superficial (SL) and deep (DL) layer neurogenesis in human. Data are from Nowakowski et al., 2018. (**B**) MiR-137 expression in VZ and oSVZ of macaque cortex during superficial layer neurogenesis. Data are from Arcila et al., 2014. (**C**) Gene ontologies of miR-137 (left) and miR-122 (right) predicted targets in human. (**D**) Endogenous expression of miR-137 (top) and miR-122 (bottom) in E15.5 mouse cortex assessed with DFRS 137 and DFRS 122, respectively (see Methods). Validation of miR-137 (DFRS 137 + miR137) and miR-122 (DFRS 122 + miR122) overexpression tools. (**E**) *In situ* hybridizations showing expression of miR-137 (left) and miR-122 (right) in mouse cortex at E14.5. Data are from Eurexpress database (http://www.eur-express.org). Data are represent ed as mean ± SEM. (E) One-way ANOVA. Biological replicates are distinguished by circles in the bar plots. *p < 0.05, **p < 10, ***p < 10^-3^, ****p < 10^-4^.

**Figure S2:**
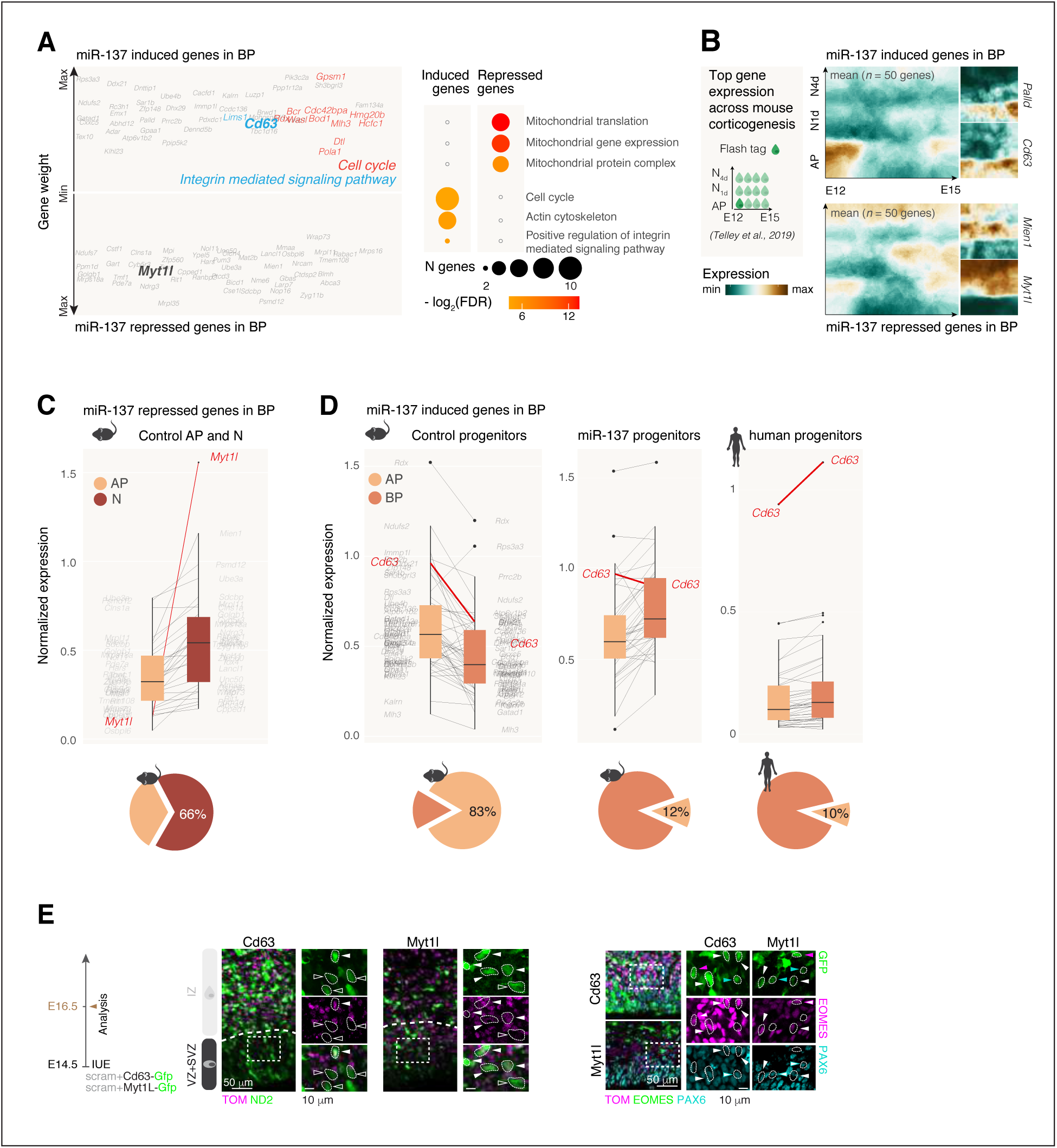
Transcriptional regulations upon miR-137 overexpression in basal progenitors. (**A**) Top 50 induced and repressed genes upon miR-137 overexpression in basal progenitors (BPs) at E15.5. Left, gene weights based on the support vector machine learning. Right, gene ontologies. (**B**) Expression of miR-137 induced and repressed genes in BPs in FlashTag labeled cells. Data are from Telley et al., 2019. (**C-D**) Expression of miR-137 repressed and induced genes in BPs in control apical progenitors (AP) and neurons (N) (**C**), and in APs and BPs from control, miR-137 and human conditions (**D**), respectively. Pie charts represent the proportion of genes expressed in AP and N (C), and in AP and BP (D). Note that miR-137 induced genes in BPs are more expressed in control APs than BPs in mouse, while in both miR-137 and human conditions they are enriched in BPs. (**E**) Representative micrographs of NeuroD2 (left) and PAX6 / EOMES (right) immunohistochemistry performed after *Cd63* and *Myt1l* overexpression. In micrographs of EOMES / PAX6 labelings: magenta arrowheads, EOMES^+^ electroporated cells; cyan arrowheads, PAX6^+^ electroporated cells, white arrowheads, EOMES^+^/PAX6^+^. Data are represented as median ± SD.

**Figure S3.**
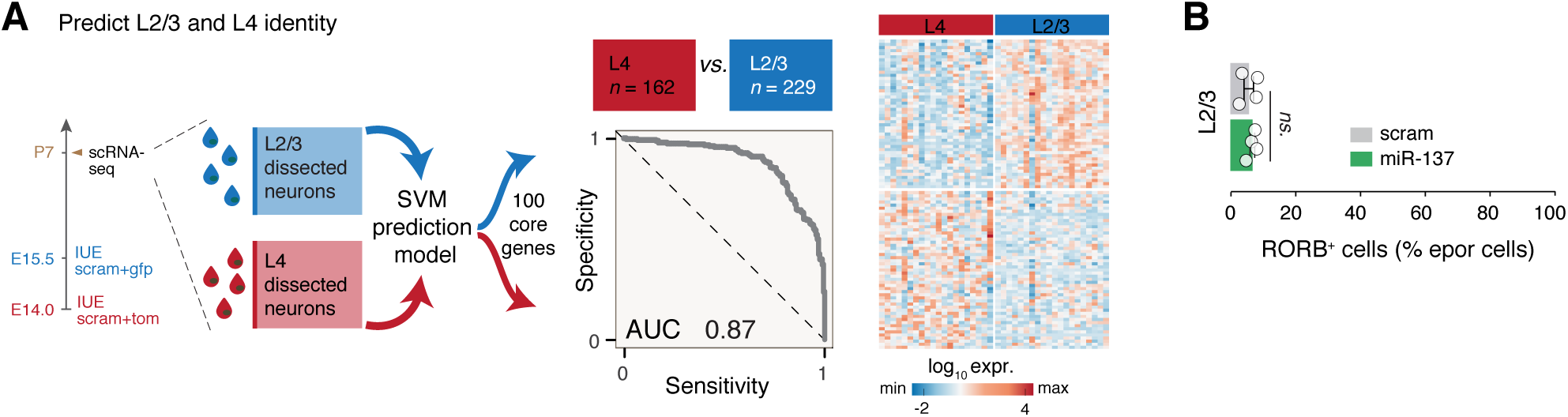
Layer 2/3 versus layer 4 molecular identity. (**A**) Left, schematic diagram of L2/3 and L4 P7 neuron isolation through birthdate-locked *in utero* electroporation (IUE). Middle, model performance using support vector machine learning approach to predict L2/3 *vs.* L4. Right, heatmap of top 100 genes which better defines L2/3 and L4 at P7. (**B**) RORB expression in L2/3 neurons in scram and miR-137 conditions. Data are represented as mean ± SEM. Biological replicates are distinguished by circles in the bar plots. Two-way ANOVA (B).

**Figure S4:**
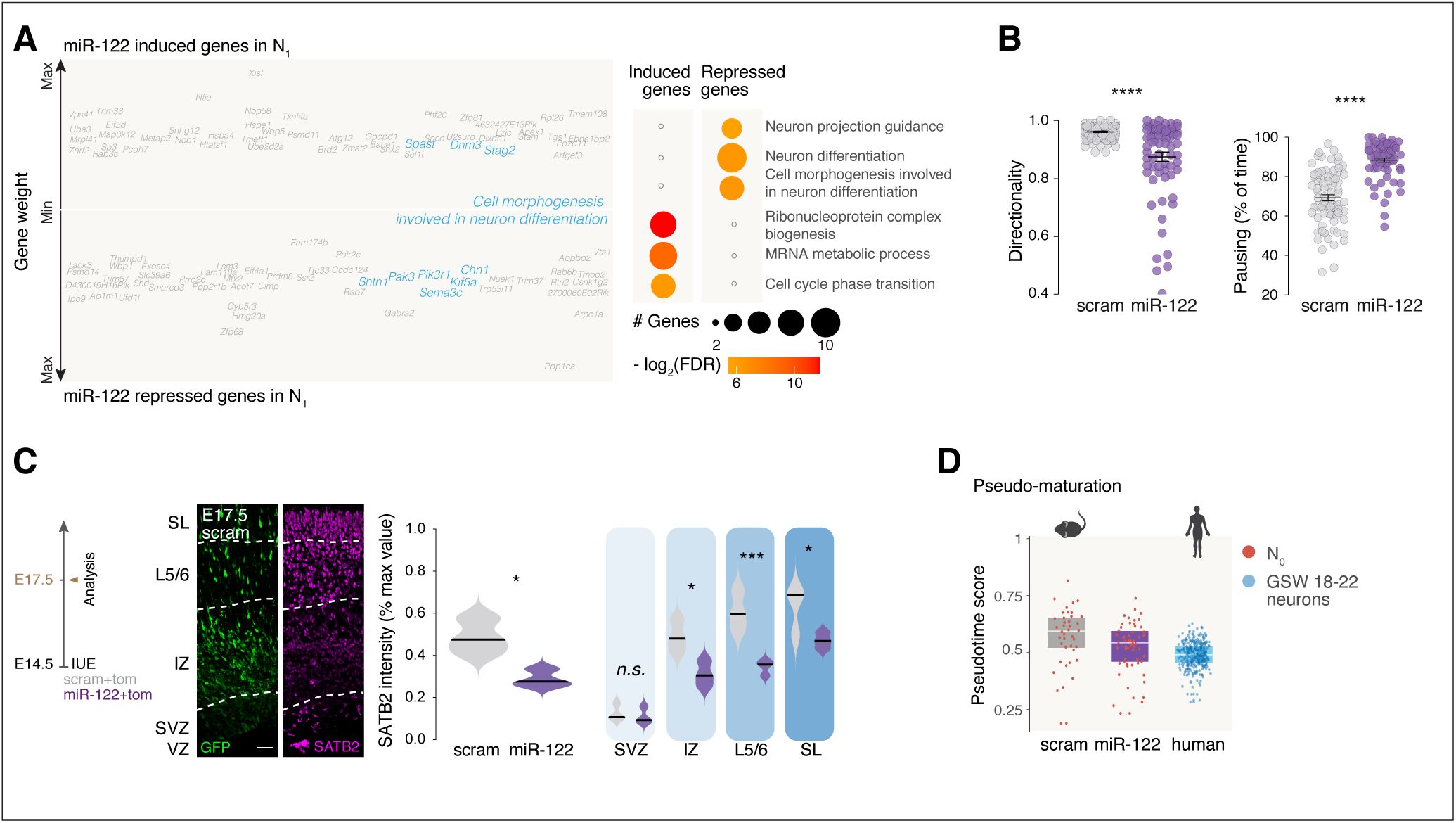
MiR-122 controls the migration and maturation of differentiating neurons. (**A**) Top 50 induced and repressed genes upon miR-122 overexpression in differentiating neurons (N_1_) at E15.5. Left, gene weights based on the support vector machine learning. Right, gene ontologies. (**B**) Directionality and pausing time of scrambled (scram) and miR-122 migrating neurons at E17.5. (**C**) Expression of SATB2, a marker for SL mature neurons, in E17.5 scram and miR-122 neurons. (**D**) Pseudotime value predictions of human superficial layer immature neurons from gestational weeks (GSW) 18-22 foetuses using the mouse model. Data are represented as mean ± SEM. Kruskal-Wallis (B); Unpaired t-Test (H, all layers); Two-way ANOVA (H, by layer). *p < 0.05, **p < 10^-2^, ***p < 10^-3^, ****p < 10^-4^. Human scRNA-seq data are from Nowakowski et al., 2017.

**Figure S5:**
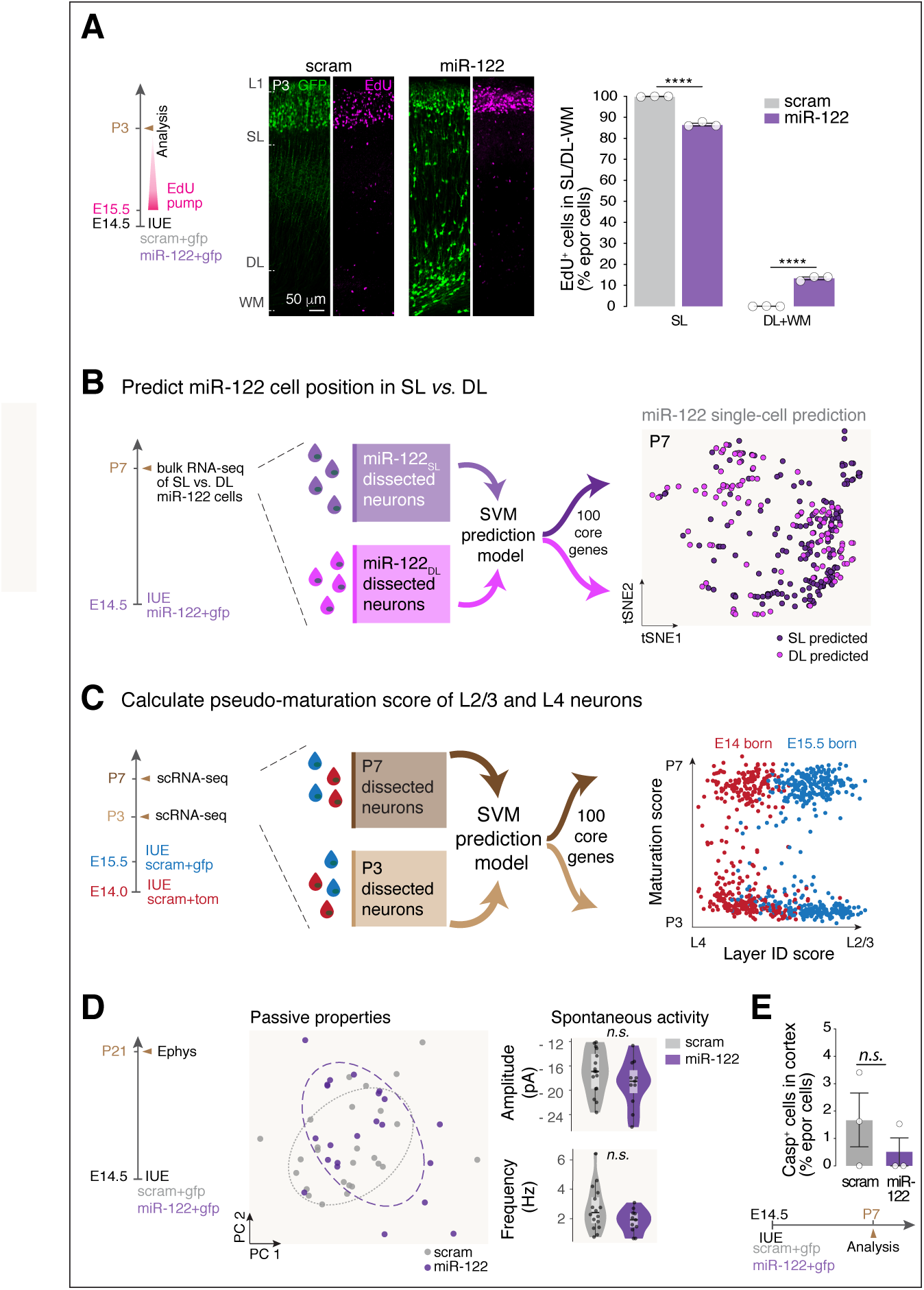
MiR-122 affects the laminar allocation and postnatal functional maturation of superficial layer cortical neurons. (**A**) Left, schematic diagram of the experiment to distinguish L2/3 neurons (born from E15.5 on) from L4 (born at E14.5) neurons after E14.5 *in utero* electroporations. Middle, EdU staining at P3. Right, Quantifications of EdU+ cells in superficial (SL) and deep (DL) layers/ white matter (WM) upon scram or miR-122 overexpression. (**B**) Left, schematic diagram of miR-122 SL and DL P7 neuron isolation after microdissection, bulk RNA-sequencing and support vector machine learning approach to predict SL *vs.* DL neuron position. Right, Prediction of miR-122 single cells using this model. (**C**) Left, schematic diagram of L2/3 and L4 P3 and P7 neuron isolation through birthdate-locked *in utero* electroporation (IUE), single-cell RNA sequencing and support vector machine learning approach to predict P3 *vs*. P7 identity (*ie*. maturation score). Right, maturation and layer identity (ID) scores of P3 and P7 E14 / E15.5 born neurons. (**D**) Passive properties and spontaneous activity recordings in L2/3 scrambled (scram) and miR-122 neurons at P21. Passive properties: Ih current, Capacitance, Input resistance, Membrane constant, Sag ratio, Spike threshold, AHP, Rheobase, Spike peak, Spike ratio, RMP, V_drop_, V_min_, V_end_, Spike delay. (**E**) CASP3 expression in cortex at P7. Quantification of CASP3+ in scram and miR-122 overexpressing cells. Data are represented as mean ± SEM (A, E) or mean ± SD (D). (A) Two-way ANOVA; D-E, Unpaired t-Test. Biological replicates and recorded cells are distinguished by circles in the bar plots (A and D, respectively). *p < 0.05, **p < 10^-2^, ***p < 10^-3^, ****p < 10^-4^.

